# Highly diverse Asgard archaea participate in organic matter degradation in coastal sediments

**DOI:** 10.1101/858530

**Authors:** Mingwei Cai, Yang Liu, Xiuran Yin, Zhichao Zhou, Michael W. Friedrich, Tim Richter-Heitmann, Rolf Nimzyk, Ajinkya Kulkarni, Xiaowen Wang, Wenjin Li, Jie Pan, Yuchun Yang, Ji-Dong Gu, Meng Li

## Abstract

Asgard is an archaeal superphylum that might hold the key to understand the origin of eukaryotes, but its diversity and ecological roles remain poorly understood. Here, we reconstructed 15 metagenomic-assembled genomes (MAGs) from coastal sediments covering most known Asgard archaea and a novel group, which is proposed as a new Asgard phylum named as the “Gerdarchaeota”. Genomic analyses predict that Gerdarchaeota are facultative anaerobes in utilizing both organic and inorganic carbon. Unlike their closest relatives Heimdallarchaeota, Gerdarchaeota have genes encoding for cellulase and enzymes involving in the tetrahydromethanopterin-based Wood–Ljungdahl pathway. Transcriptomic evidence showed that all known Asgard archaea are capable of degrading organic matter, including peptides, amino acids and fatty acids, in different ecological niches in sediments. Overall, this study broadens the diversity of the mysterious Asgard archaea and provides evidence for their ecological roles in coastal sediments.

## Introduction

Asgard archaea, proposed as a new archaeal superphylum, are currently composed of five phyla, i.e., Lokiarchaeota^1^, Thorarchaeota^2^, Odinarchaeota^3^, Heimdallarchaeota^3^, and Helarchaeota^4^, some of which encompasses lineages formerly named Marine Benthic Group B (MBG-B)^5^, Deep-Sea Archaeal Group (DSAG)^6^, Ancient Archaeal Group (AAG)^7^, and Marine Hydrothermal Vent Group (MHVG)^7, 8^. Since Asgard archaea contain abundant eukaryotic signature proteins (ESPs) and form a monophyletic group with eukaryotes in the phylogenetic tree inferred from 55 concatenated archaeo-eukaryotic ribosomal proteins, they are regarded as the closest relatives of Eukarya and have attracted increasing research interests^1, 3, 9^.

Metabolic potentials of several Asgard phyla have been predicted based upon their genomic inventories: hydrogen dependency^10^ in Lokiarchaeota (which were originally described in deep-sea sediments), mixotrophy^11^ (i.e., using both inorganic and organic carbon for growth) and acetogenesis^2^ by Thorarchaeota from estuary sediments, metabolization of halogenated organic compounds^12^ by Loki- and Thorarchaeota, phototrophy^13, 14^ in Heimdallarchaeota from coastal sediment and hydrothermal vent samples, anaerobic hydrocarbon oxidation in hydrothermal deep-sea sediment by Helarchaeota^4^, and nitrogen and sulfur cycling^15^ in all Asgard archaeal phyla. Metatranscriptomics revealed a few transcripts encoding NiFe hydrogeneases from deep-sea Lokiarchaeota^16^. Since no comprehensive information about the *in situ* activity of these Asgard archaea has been compiled so far, our understanding of these phylogenetically and evolutionally important archaea is based on prediction, and thus, severely limited.

Coastal environments, e.g., mangroves, salt marshes and seagrass beds, are known sinks of blue carbon^17^. Although these vegetated coastal ecosystems make up less than 0.5% of the seabed, they hold ∼50% of organic carbon of the surface marine sediments globally^17-19^. Here, we reconstructed 15 Asgard metagenomic-assembled genomes (MAGs) from diverse coastal sediments and analysed them together with transcriptomes from mangrove sediments to clarify the ecological roles of the different Asgard clades in these important environments. Based on phylogenetic analysis of this dataset, we propose a novel Asgard phylum, the “Gerdarchaeota”. Additionally, we recruited all publicly available 16S rRNA sequences to see whether or not distinct Asgard lineages show distinct habitat preferences. Our findings substantially extend our knowledge of the lifestyles of these mysterious archaea in coastal sediments and their ecological roles on carbon cycling.

## Results and Discussion

### Mining Asgard archaea genomes lead to the proposal of a new Asgard phylum

Sediments from several coastal sites (mangrove, mudflat and seagrass bed) were collected for deep metagenomic and metatranscriptomic sequencing (totally 2.3 Tbp, table S1). By combining individual samples from the same site but from different depth layers for assembly and binning, we recovered 15 Asgard MAGs with completeness of >80% through hybrid binning strategies (MetaBAT^20^ in combination with Das Tool^21^) and manual bin refinement. Based on a phylogenetic analysis with a concatenated set of 122 archaeal-specific marker genes, we found that eight of the MAGs belong to known Asgard lineages, covering almost all phyla (except Odinarchaeota), i.e., Helarchaeota (1 MAG), Lokiarchaeota (2), Thorarchaeota (3), Heimdallarchaeota-AAG (1), and Heimdallarchaeota-MHVG (1) (Fig. 1A and table S2).

**Fig. 1.**
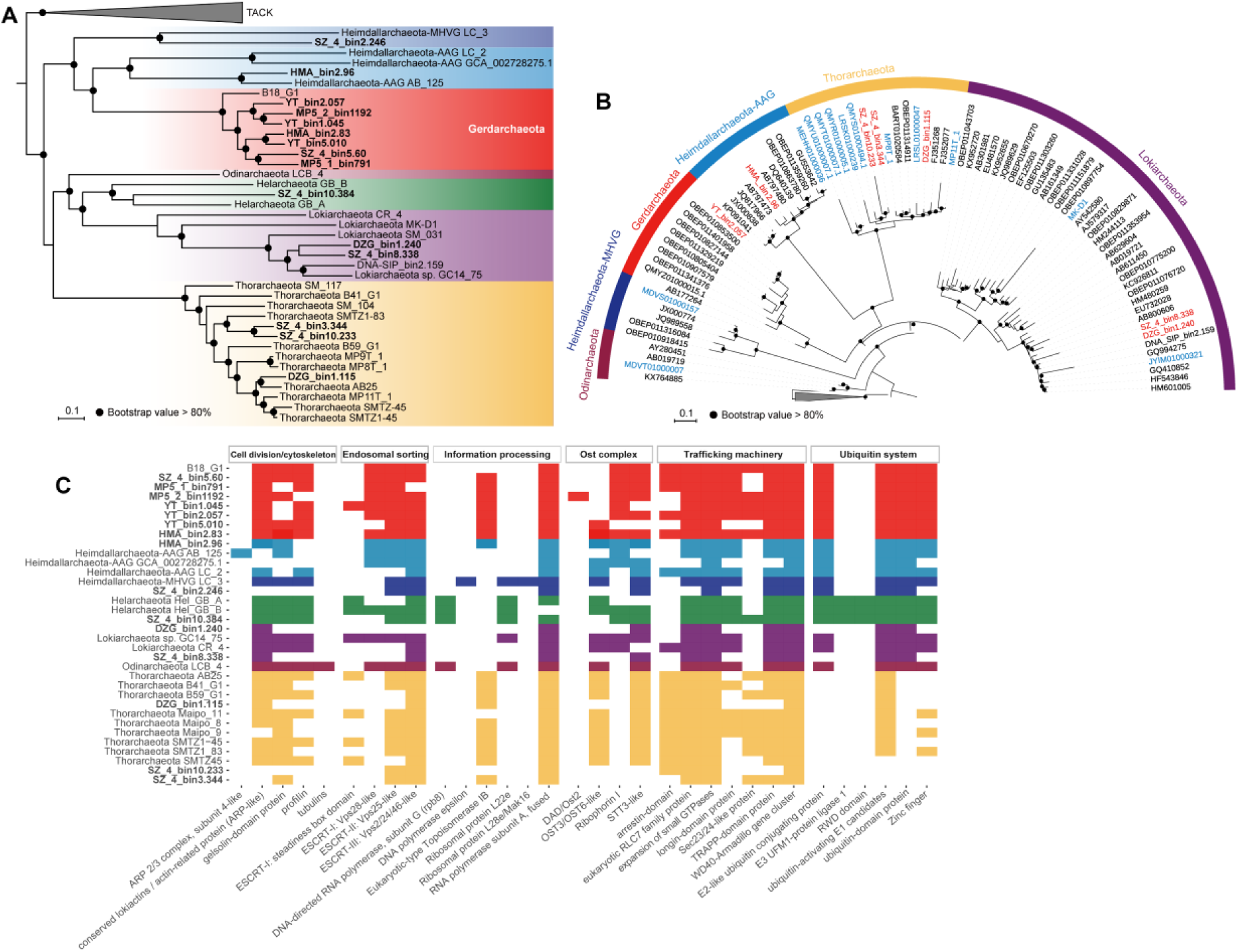
Phylogenetic positions and ESPs of Asgard archaea reconstructed from coastal sediments. (**A**), Maximum likelihood tree of Gerdarchaeota MAGs built on a concatenated alignment of 122 archaeal marker genes. The tree was inferred with LG+F+R10 mixture mode in IQ-TREE and rooted with DPANN and Euryarchaeota. Asgard archaea MAGs obtained in this study are marked in bold face. (**B**), Phylogenetic position of Asgard archaeal 16S rRNA genes. Red colour represents 16S rRNA genes from newly discovered Asgard MAGs, and blue colour represents sequences from reference ones. The tree was built using the IQ-TREE software with GTR+I+G4 mixture mode and rooted with Crenarchaeota. (**C**), ESPs identified in Gerdarchaeota and other Asgard archaea. Asgard archaea MAGs obtained in this study are marked bold. Colours indicate phylum-level assignment (see 3A).

The remaining 7 MAGs clustered with an unclassified MAG (B18_G1) and formed a monophyletic group in phylogenetic tree of concatenated 122 archaeal marker genes (Fig. 1A). The amino acid identity (AAI, 43% to 50% compared to other Asgard archaea), pan-genome analysis, and non-metric multidimensional scaling (NMDS) ordination agreed on assigning phylum level identity to this new clade, (figs. S1, S2 and S3). Likewise, phylogenetic analysis of a 16S rRNA (1152 bp) gene found in of those seven MAGs (YT_re_metabat2_2.057) showed that it formed a new monophyletic branch (Fig. 1B) with an identity below 74% to 16S rRNA genes of other Asgard archaeal phyla (table S3). Thus, we propose this lineage as a new phylum named Gerdarchaeota, after Gerd, the Norse goddess of fertile soil, because these Asgard archaea genomes were obtained from organic-rich coastal environments^22^, such as mangrove, mudflat and seagrass (table S1).

The presence of eukaryotic signature proteins (ESPs) is a characteristic feature of Asgard (Fig. 1C and table S4), and we consistently found that their phylogenies support the discrete position of Gerdarchaeota in the Asgard lineage (fig. S4). Homologs encoding eukaryotic-type topoisomerase IB and fused RNA polymerase subunit A were identified in Gerdarchaeota, while neither genes for eukarya-specific DNA polymerase epsilon and DNA-directed RNA polymerase subunit G were detected (Fig. 1C, and table S5). Gerdarchaeotal topoisomerase IB is clustered into a sister group of Thorarchaeota and the two Asgard sequences together were monophyletic with high supporting values (fig. S4A). The fused RNA polymerase A genes of Gerdarchaeota are transcriptionally expressed (table S5) and clustered into a sister group of Heimdallarchaeota, and those of Helarchaeota branch with Loki- and Thorarchaeota (fig. S4B). In both cases, the eukaryotic genes do not cluster with the Asgard homologs. Besides, Gerdarchaeota also comprise expressed homologues of ribophorin I and STT3 subunit and lack OST3/OST6 homologues in most genomes except YT_bin5.010. The phylogenetic tree of ribophorin I showed that Gerdarchaeota is monophyletic and branches with Heimdallarchaeota, and the eukaryotes branch within the cluster of Thor-, Hel- and Lokiarchaeota (fig. S4C). In addition to the reported ESPs, we identified DAD/OST2 homologues within Gerdarchaeotal MAGs, which is a component of the N-ologosaccharyl transferase for the N-linked glycosylation^23^.

### Metabolic potential of the new phylum Gerdarchaeota

Gerdarchaeota harbour the gene set for oxidative phosphorylation, including V/A-type ATPs, succinate dehydrogenase, NADH-quinone oxidoreductase, the key enzyme cytochrome *c* oxidase (subunits I, II, and III) (Fig. 2, fig. S5, and tables S6 and S7), and enzymes for the non-typical cytochrome *bc1* complex (i.e., SoxL)^24^ but lacking the genes for other respiratory complex III (e.g.,SoxN and CbsA), suggesting that Gerdarchaeota most likely perform aerobic respiration. Gerdarchaeotal cytochrome *c* oxidases are phylogenetically separated into two lineages (fig. S6), one of which clusters closely with the facultative anaerobic Crenarchaeotal *Acidianus brierleyi* ^25, 26^ and the other one groups with the bacterial *Synechocystis* sp., which is capable of aerobic respiration in the dark^27^. Meanwhile, Gerdarchaeotal MAGs harbour genes encoding heliorhodopsins (fig. S7), which might sense light in the top layers of sediment^13, 14^. A complete tricarboxylic acid cycle (e.g., citrate synthase and malate dehydrogenase, Fig. 2) further supports aerobic respiration as important dissimilatory pathway. Besides, Gerdarchaeota are equipped with enzymes for the conversion of AsO_4_^3-^ to (CH_3_)_n_-As, thus they could remove the toxic As(V) and As(III), which can disrupt oxidative phosphorylation and inhibit respiratory enzymes^28^.

**Fig. 2.**
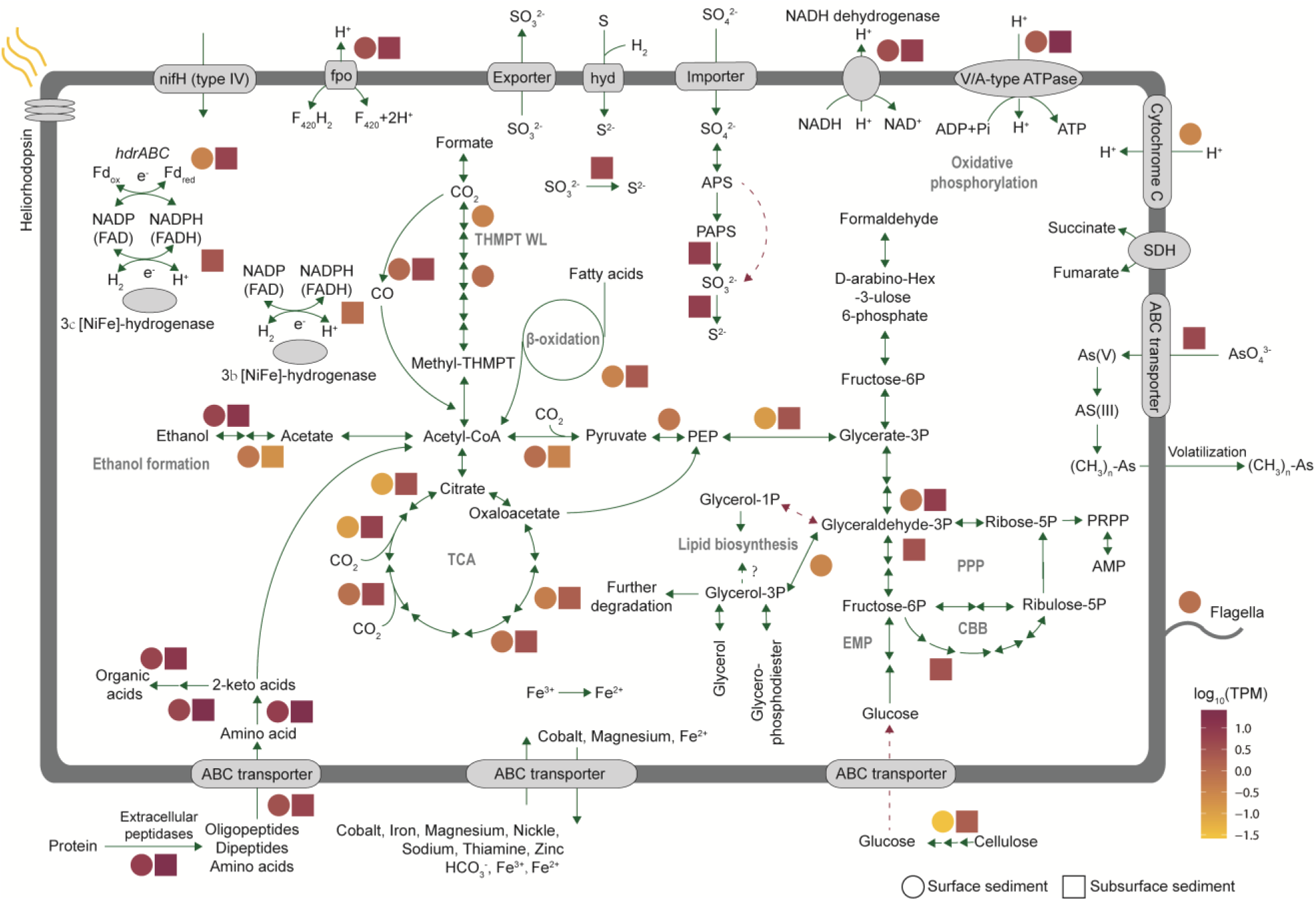
Key potential metabolic pathways of Gerdarchaeota. Solid lines represent pathway steps present in 8 MAGs assigned to Gerdarchaeota. Dashed lines represent pathway steps not identified. Circles represent transcripts from surface sediments and rectangles represents transcripts from subsurface samples. The relative abundance of the transcripts (Transcripts Per Million reads, TPM) for each gene is marked with different colours. Pathway abbreviations: PPP, pentose phosphate pathway (archaea); CBB, Calvin–Benson–Bassham cycle; EMP, Embden–Meyerhof–Parnas pathway, THMPT WL, tetrahydromethanopterin-dependent Wood–Ljungdahl pathway; TCA cycle, tricarboxylic acid cycle. Detailed metabolic information for the MAGs is available in fig. S5, tables S6, and S7.

Within Gerdarchaeota MAGs, complete gene sets for acetogenesis pathways (e.g., acetyl-CoA hydrolase) are present (Fig. 2), implying the ability of these archaea to perform fermentation in the absence of oxygen. We further identified all subunits of [NiFe]-hydrogenase heterodisulfide reductase *hdrABC*. Since this community does not contain the key enzymes *mcrABC* for methanogenesis, *hdrABC* might function to accept electrons and reduce oxidized ferredoxin instead of a heterodisulfide anaerobically^29-32^. Additionally, we identified homologues for ferric reductase but their capability to use Fe(III) as electron acceptors remained open. Notably, the canonical nitrate reductase (previously identified in Heimdallarchaeota^32^) was not detected in Gerdarchaeota MAGs.

Gerdarchaeota appear to use diverse organic compounds (e.g., formaldehyde, amino acid, peptide, lipid and ethanol) as electron donors (Fig. 2). For genes encoding peptidase, serine peptidases dominated (∼ 44.1% of total peptidase, fig. S8A). We also identified abundant genes for cellulose degradation (e.g., AA3 and GT2, fig. S8B and table S8), which may be further degraded through the Embden–Meyerhof–Parnas (EMP) pathway. Different from the facultative anaerobic relatives Heimdallarchaeota, Gerdarchaeota contain the complete genes for tetrahydromethanopterin Wood-Ljungdahl (THMPT_WL) pathway and the key enzyme acetyl-CoA decarbonylase/synthase (Fig. 2). The presence of the genes for groups 3b and 3c [NiFe]-hydrogenases (fig. S9) implies that these archaea may grow lithoautotrophically using H_2_ as electron donor as also reported for Lokiarchaeota^10^. Meanwhile, we identified other CO_2_ assimilation pathways in Gerdarchaeota MAGs. For example, Gerdarchaeota are equipped to fix carbon via the complete reductive citric acid cycle (e.g., 2-oxoglutarate oxidoreductase and isocitrate/isopropylmalate dehydrogenase) and Calvin–Benson–Bassham (CBB) cycle, and the pyruvate dehydrogenase, which is responsible for the generation of pyruvate from acetyl-CoA and CO_2_, underpin the importance of inorganic carbon for biomass synthesis^33^. Different from other Asgard phyla, we did not find type III or type IV ribulose 1,5-bisphosphate carboxylase (RuBisCO) (fig. S10), which might function in the nucleotide salvage pathway^11, 21, 26, 27.^

Most reported Asgard genomes contain genes for Glycerol-1-phosphate dehydrogenase (G1PDH) responsible for ether-bound phospholipids synthesis^12, 33^. A recent study reported a co-existence of enzymes for both ether- (G1PDH) and ester-bound (bacterial/eukarya type, glycerol-3-phosphate dehydrogenase (G3PDH)) phospholipid synthesis in some Lokiarchaeotal genomes, providing a hint that Asgard archaea might produce chimeric lipids^12^. Intriguingly, Gerdarchaeota MAGs lack the key enzyme G1PDH for archaeal lipid biosynthesis, but contain the bacterial/eukarya-type G3PDH for synthesis of bona fide bacterial lipids (Fig. 2). This finding indicated that Gerdarchaeota might have evolved a bacterial-like membrane predates the eukaryogenesis. Further investigation suggests that the Asgard archaeal G3PDH is synthesized by the *glpA* (responsible for the reverse reaction catalyzed by GpsA) instead of GpsA, which means that G3PDH may participate in organic carbon degradation rather than lipid synthesis^34, 35^, leaving the mechanism of lipid synthesis to be further explored.

### Transcriptomic activities of Asgard archaea in different niches of coastal sediments

Gene expression patterns derived from metatranscriptomic analysis have been used in a number of recent studies to deduce active microbial processes in marine sediments, especially in the deep sea^36-38^. This technique might be constrained by the largely unknown pool size of mRNAs maintained in endospores and dormant cells^39^, although the former seems to be irrelevant for Asgard archaea. Through recruiting 16S rDNA/RNA sequences (n=10,448, read length >600bp) belonging to Asgard archaea from public databases, we found that all Asgard phyla were transcriptionally expressed and diversely distributed (with ∼92% of Asgard OTUs originated from sediment samples, fig. S11 and table S9). Thus, to better uncover their activities, we used 818,479 transcripts (mRNA) belonging to Asgard archaea including Loki-, Thor-, Hel-, Heimdall- and Gerdarchaeota from mangrove sediments to elucidate their ecological roles in coastal sediments.

Organic carbons in coastal sediments are mainly composed of carbohydrates, amino acids, and lipids^22, 40^; here, amino acids are the most dominant component of organics in mangrove sediments^41^. Accordingly, we detected high expression levels of genes encoding extracellular peptidases, ABC transporter, and the enzyme sets for the conversion of amino acids to acetyl-CoA in both surface and subsurface coastal sediments, implying that Asgard archaea might be essential participants in the degradation of these substrates (Fig. 2, fig. S5, and tables S6, S7 and S10). This feature is supported by the high proportion of peptidases in Asgard archaea MAGs (4.1–6.3 % of the functional genes, fig. S8A). We also detected transcripts for ethanol metabolism, suggesting that ethanol might be another substrate or product. Although glucose accounts for 6–18% of the total organic carbon in mangrove sediments^41^, it might be not the first nutritional choice for Asgard archaea because no transcript for the key enzyme glucokinase was detected.

Due to its higher energetic efficiency compared to fermentative processes^43^, aerobic respiration contributes to 50% or more of the total organic matter decomposition in the offshore marine sediments. The expressed gene set for aerobic respiration in Gerdarchaeota, Heimdallarchaeota-AAG and Heimdallarchaeota-MHVG, including the key transcript of cytochrome *c* oxidase (belonging to Gerdarchaeota) indicates that these Asgard archaea might participate aerobically in organic matter degradation in surface sediments (Fig. 3). Although co-existing within the same depth layers, Asgard archaea might play phylum specific ecological roles. For example, unlike Heimdallarchaeota-AAG and Heimdallarchaeota-MHVG, Gerdarchaeota contain and expressed genes for autotrophy and cellulose degradation, but lack the complete gene set for glucose degradation (Fig. 3). Like Gerdarchaeota, other Asgard archaea have the potential to perform anaerobic metabolisms (e.g., acetogenesis) under anoxic conditions (subsurface layers) ^42^.Notably, Helarchaeota-like *mcrA* gene transcripts found in unbinned scaffolds (e.g., SZ_4_scaffold_203331_2, fig. S12) highlight the involvement of Helarchaeota in alkane oxidation in coastal sediments, in which ethane and butane might originate from oil-gas seepage or human activities^43^, are preferentially used as revealed by molecular modelling and dynamics studies (fig. S13 and Supplementary Results).

**Fig. 3.**
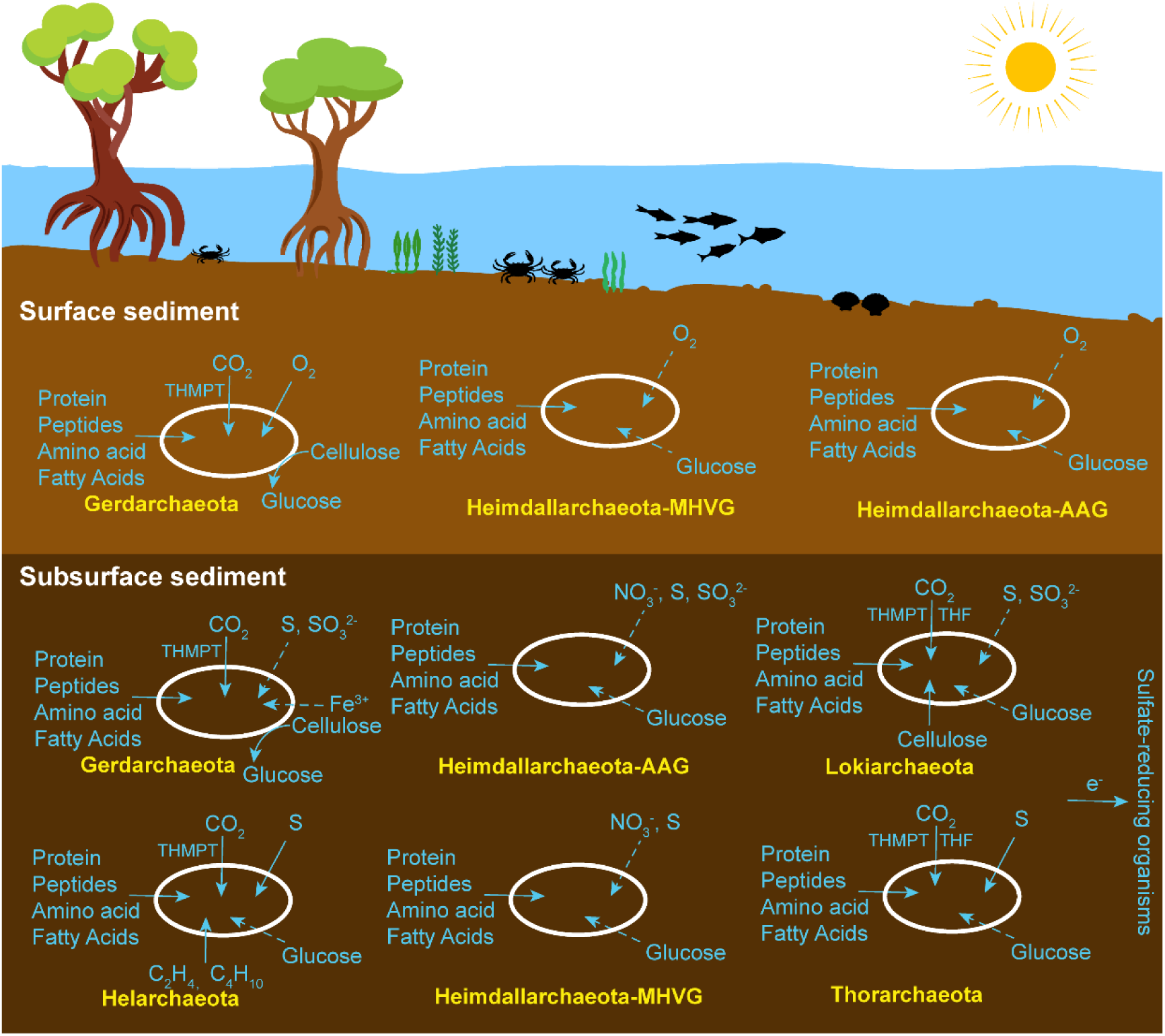
Ecological niches of Asgard archaea in coastal sediments. Dashed lines represent pathways with no transcript for the key genes. Detailed information is available in fig. S5, tables S6, and S7.

Previous studies suggested that Asgard archaea could account for up to 50% of the total prokaryotes in some marine sediments^44^; and attribute to 40% of the total archaeal sequences in coastal sediments^44, 45^. The discovery of novel lineages, as well as the discovery of co-occuring diverse lineages in one vertical biosphere in this study, may elevate their relative abundance and ecological roles in natural environments. Therefore, we propose that Asgard archaea might be essential archaeal lineages for organic carbon degradation in coastal sediments, similar to previously proposed roles for Bathyarchaeota^45^ and Thermoprofundales^46^, which also support the notion that Asgard archaea are critical participants for organic matter utilization in coastal sediments^44, 47^.

## Conclusion

Currently, the research of the diversity and ecology of Asgard archaea is still in its infancy. Contrasting previous studies that Asgard archaeal MAGs covered not more than three phyla^3, 4, 14^, our study features a wider taxonomic breadth, as we obtained almost all known Asgard phyla in the coastal sediments, and additionally found a new Asgard archaea phylum. Metabolic comparison and transcriptomic evidence suggest divergent ecological roles and niches for different Asgard phyla but they shared key transcripts involved in the degradation of specific compartments of organic matter (e.g., peptides and amino acids). Thus, considering their high relative abundance^44, 45^ and ubiquitous distribution in the coastal area, we infer that Asgard archaea are important players for organic matter utilization^45, 48^. However, their contribution to the coastal sediment carbon budget remains to be further examined. Overall, the metabolic features, transcript evidence, and their global distribution imply that Asgard archaea are essential players in carbon cycling of coastal sediments.

## Materials and Methods

### Sediment sample collection and processing

Samples for metagenome analysis were collected from the coastal sediment (i.e., mangrove, mudflat and seagrass sediments) of China and Helgoland coastal mud area during the RV HEINCKE cruise HE443 (table S1). They were sampled using custom corers, sealed in plastic bags in duplicates, stored in sampling box with ice bags, and transported to the lab within 4 hours. The physiochemical parameters of the samples were determined as previously described^49^. Samples for RNA extraction were preserved in RNALater (Ambion, Life Technologies). For each sample, 10 g of sediment each was used for DNA and RNA isolation with the PowerSoil DNA Isolation Kit (MO BIO) and RNA Powersoil™ Total RNA Isolation Kit (QIAGEN), respectively. The rRNA genes were removed from the total RNA using the Ribo-Zero rRNA removal kit (Illumina, Inc., San Diego, CA, USA) and the remaining mRNA was reverse-transcribed. DNA and cDNA were sequenced using an Illumina HiSeq sequencer (Illumina) with 150-bp paired-end reads at BerryGenomics (Beijing, China). Metatranscriptomic reads were quality-trimmed using Sickle (version 1.33)^50^ with quality score ≥25, and the potential rRNA reads were removed using SortMeRNA (version 2.0)^51^ against both the SILVA 132 database and the default databases (E-value cutoff ≤1e-5).

### Metagenomic assembly, genome binning and gene annotation

Raw metagenomic DNA reads of the coastal sediments were dereplicated (identical reads) and trimmed using Sickle (version 1.33)^50^ with the option “-q 25”. Paired-end Illumina reads for each sample were assembled *de novo* using IDBA-UD (version 1.1.1)^52^ with the parameters “-mink 65, -maxk 145, -steps 10”. Scaffolds were binned into genomic bins using a combination of MetaBAT^20^ and Das Tool^21^. Briefly, twelve sets of parameters were set for MetaBAT binning, and Das Tool was further applied to obtain an optimized, non-redundant set of bins. To improve the quality of the bins (e.g., scaffold length and bin completeness), each Asgard-related bin was remapped with the short-read mapper BWA and re-assembled using SPAdes (version 3.0.0)^53^ or IDBA-UD (version 1.1.1)^52^, followed by MetaBAT and Das Tool binning. Asgard MAGs with high contamination were further refined with Anvi’o software (version 2.2.2)^54^. The completeness, contamination and strain heterogeneity of the genomic bins were estimated by CheckM (version 1.0.7) software^55^. Anvi’o software (version 2.2.2)^54^ was applied for pan-genome analysis of Asgard MAGs with the option “--min-occurrence 3”.

Protein-coding regions were predicted using Prodigal (version 2.6.3) with the “-p meta” option^56^. The KEGG server (BlastKOALA)^57^, eggNOG-mapper^58^, InterProScan tool (V60)^59^, and BLASTp vs. NCBI-nr database searched on December 2017 (E-value cutoff ≤1e-5) were used to annotate the protein-coding regions. Archaeal peptidases were predicted against the MEROPS database^60^, and the extracellular peptidases were further identified using PRED-SIGNAL^61^ and PSORTb^62^ (table S11).

### Phylogenetic analyses of Asgard MAGs

The 16S rRNA gene sequences and a concatenated set of 122 archaeal-specific conserved marker genes^63^ were used for phylogenetic analyses of Asgard archaea. Ribosomal RNA genes in the Asgard MAGs were extracted by Barrnap (version 0.3, http://www.vicbioinformatics.com/software.barrnap.html). An updated 16S rRNA gene sequence dataset from reference papers^64,65^ with genome-based 16S rRNA genes were aligned using SINA (version 1.2.11)^66^. The 16S rRNA gene sequences maximum-likelihood tree was built with IQ-TREE (version 1.6.1)^67^ using the GTR+I+G4 mixture model (recommended by the “TESTONLY” model), with option “-bb 1000”. Marker genes for protein tree were identified using hidden Markov models (HMMs) and were aligned separately using hmmalign from HMMER3^68^ with default parameters. The 122 archaeal marker genes were identified using hidden Markov models. Each protein was individually aligned using hmmalign^69^. The concatenated alignment was trimmed by BMGE with flags “-t AA -m BLOSUM30”^70^. Then, maximum-likelihood trees were built using IQ-TREE with the best-fit model of “LG+F+R10” followed by extended model selection with FreeRate heterogeneity and 1000 times ultrafast bootstrapping. The final tree was rooted with the DPANN superphylum and Euryarchaeota.

### Metabolic pathway construction

Potential metabolic pathways were reconstructed based on the predicted annotations and the reference pathways depicted in KEGG and MetaCyc^71^. Metatranscriptome data from mangrove and mudflat sediments of Shenzhen Bay (table S1) were analyzed to clarify the transcriptomic activity of Asgard archaea. The abundance of transcripts for each gene was determined by mapping all non-rRNA transcripts to predicted genes using BWA with default setting^36, 72^. Normalized expression was expressed in transcript per million units (TPM), followed by normalization by genome number of each phylum.

### ESP identification

As predicted by prodigal (v2.6.3) with default parameters, genes of Gerdarchaeota were searched against InterPro and eggNOG databases to gain the IPRs and arCOGs. The list of those annotations in Zaremba-Niedzwiedzka et al (2017) ^3^ was searched for in the Gerdarchaeota bins. We also manually inspected the IPRs and arCOGs only present in Gerdarchaeota. MAFFT-linsi and trimAl (-gappyout) were used to align and trim the protein sequences. IQ-TREE (version 1.6.1)^67^ was used to infer phylogeny of under best-fit models with 1000 ultrafast bootstraps with SH-aLRT test values.

## General

We thank Dr. Nidhi Singh for her suggestions in molecular modeling. We thank the captain, crew and scientists of R/V HEINCKE expeditions HE443

## Author contributions

M.L., M.C. and Y.L. conceived this study. M.C. analyzed the 16S rRNA data, metagenomic data and metatranscriptomic data. Y.L. collected samples and analyzed the metagenomic data. X.Y., M.W.F., T.R.H., R.N., and A.K. provided metagenomic data. Z.Z. provided support for diversity analysis. M.C., Y.C.Y., J.P. and Z.Z. prepared the DNA and cDNA for sequencing. W.L. and X.W. analyzed MCR complex protein structure and simulated the binding substrates. M.C., Y.L., and M.L. wrote, and all authors edited and approved the manuscript.

## Data and materials availability

Archaeal 16S rRNA gene sequences were retrieved from NCBI database, SILVA SSU r132 database, and a reference paper as described in Supplementary Materials and Methods. Public Asgard MAGs were from NCBI database and MG-RAST. The newly obtained Asgard MAGs and metatranscriptomic data are available in NCBI database under the project PRJNA495098 and PRJNA360036.

## Supplementary Materials

Supplementary methods and results

figures S1 to S16

tables S1 to S12

## References and Notes

1. Spang, A. et al. Complex archaea that bridge the gap between prokaryotes and eukaryotes. Nature 521, 173–179 (2015).

2. Seitz, K. W. et al. Genomic reconstruction of a novel, deeply branched sediment archaeal phylum with pathways for acetogenesis and sulfur reduction. ISME J. 10, 1696 (2016).

3. Zaremba-Niedzwiedzka, K. et al. Asgard archaea illuminate the origin of eukaryotic cellular complexity. Nature 541, 353–358 (2017).

4. Seitz, K. W. et al. Asgard archaea capable of anaerobic hydrocarbon cycling. Nat. Commun. 10, 1822 (2019).

5. Vetriani, C. et al. Population structure and phylogenetic characterization of marine benthic archaea in deep-sea sediments. Appl. Environ. Microbiol. 65, 4375–4384 (1999).

6. Inagaki, F. et al. Archaeology of Archaea: geomicrobiological record of Pleistocene thermal events concealed in a deep-sea subseafloor environment. Extremophiles 5, 385–392 (2001).

7. Takai, K. & Horikoshi, K. Genetic diversity of archaea in deep-sea hydrothermal vent environments. Genetics 152, 1285–1297 (1999).

8. Inagaki, F. et al. Microbial communities associated with geological horizons in coastal subseafloor sediments from the Sea of Okhotsk. Appl. Environ. Microbiol. 69, 7224–7235 (2003).

9. Zhou, Z., Liu, Y., Li, M. & Gu, J. D. Two or three domains: a new view of tree of life in the genomics era. Appl. Microbiol. Biot. 102, 3049–3058 (2018).

10. Sousa, F. L. et al. Lokiarchaeon is hydrogen dependent. Nat. Microbiol. 1, 16034 (2016).

11. Liu, Y. et al. Comparative genomic inference suggests mixotrophic lifestyle for Thorarchaeota. ISME J. 12, 1021–1031 (2018).

12. Manoharan, L. et al. Metagenomes from Coastal Marine Sediments Give Insights into the Ecological Role and Cellular Features of Loki- and Thorarchaeota. mBio 10, e02039–19 (2019).

13. Pushkarev, A. et al. A distinct abundant group of microbial rhodopsins discovered using functional metagenomics. Nature 558, 595–599 (2018).

14. Bulzu, P.-A. et al. Casting light on Asgardarchaeota metabolism in a sunlit microoxic niche. Nat. Microbiol. 4, 1129–1137 (2019).

15. MacLeod, F. et al. Asgard archaea: Diversity, function, and evolutionary implications in a range of microbiomes. AIMS Microbiol. 5, 48–61 (2019).

16. Huang, J.-M., Baker, B. J., Li, J.-T. & Wang, Y. New microbial lineages capable of carbon fixation and nutrient cycling in deep-sea sediments of the northern South China Sea. Appl. Environ. Microbiol. 85, e00523–19 (2019).

17. Mcleod, E. et al. A blueprint for blue carbon: toward an improved understanding of the role of vegetated coastal habitats in sequestering CO2. Front. Ecol. Environ. 9, 552–560 (2011).

18. Breithaupt, J. L. et al. Organic carbon burial rates in mangrove sediments: Strengthening the global budget. Global Biogeochem. Cy. 26, (2012).

19. Kennedy, H. et al. Seagrass sediments as a global carbon sink: Isotopic constraints. Global Biogeochem. Cy. 24, (2010).

20. Kang, D. D., Froula, J., Egan, R. & Wang, Z. MetaBAT, an efficient tool for accurately reconstructing single genomes from complex microbial communities. PeerJ 3, e1165 (2015).

21. Sieber, C. M. et al. Recovery of genomes from metagenomes via a dereplication, aggregation and scoring strategy. Nat. Microbiol. 3, 836–843 (2018).

22. Miyatake, T., MacGregor, B. J. & Boschker, H. T. Depth-related differences in organic substrate utilization by major microbial groups in intertidal marine sediment. Appl. Environ. Microbiol. 79, 389–392 (2013).

23. Silberstein, S., Collins, P. G., Kelleher, D. J. & Gilmore, R. The essential OST2 gene encodes the 16-kD subunit of the yeast oligosaccharyltransferase, a highly conserved protein expressed in diverse eukaryotic organisms. J. Cell Biol. 131, 371–383 (1995).

24. Marreiros, B. C. et al. Exploring membrane respiratory chains. BBA-Bioenergetics 1857, 1039–1067 (2016).

25. Segerer, A., Neuner, A., Kristjansson, J. K. & Stetter, K. O. Acidianus infernus gen. nov., sp. nov., and Acidianus brierleyi comb. nov.: facultatively aerobic, extremely acidophilic thermophilic sulfur-metabolizing archaebacteria. Int. J. Syst. Evol. Micr. 36, 559–564 (1986).

26. Konishi, Y., Asai, S., Tokushige, M. & Suzuki, T. Kinetics of the bioleaching of chalcopyrite concentrate by acidophilic thermophile Acidianus brierleyi. Biotechnol. Prog. 15, 681–688 (1999).

27. De Rosa, E. et al. [NiFe]-hydrogenase is essential for cyanobacterium Synechocystis sp. PCC 6803 aerobic growth in the dark. Sci. Rep. 5, 12424 (2015).

28. Thiel, V. Archaea. Encyclopedia of Geobiology, 64–69 (2011).

29. Greening, C. et al. Genomic and metagenomic surveys of hydrogenase distribution indicate H2 is a widely utilised energy source for microbial growth and survival. ISME J. 10, 761–777 (2016).

30. Peters, J. W. et al. [FeFe]- and [NiFe]-hydrogenase diversity, mechanism, and maturation. BBA-Mol. Cell Res. 1853, 1350–1369 (2015).

31. Hua, Z.-S. et al. Genomic inference of the metabolism and evolution of the archaeal phylum Aigarchaeota. Nat. Commun. 9, 2832 (2018).

32. Spang, A. et al. Proposal of the reverse flow model for the origin of the eukaryotic cell based on comparative analyses of Asgard archaeal metabolism. Nat. Microbiol., 1138–1148 (2019).

33. Yin, X. et al. CO2 conversion to methane and biomass in obligate methylotrophic methanogens in marine sediments. ISME J. 13, 2107–2119 (2019).

34. Yokobori, S.-i., Nakajima, Y., Akanuma, S. & Yamagishi, A. Birth of archaeal cells: molecular phylogenetic analyses of G1P dehydrogenase, G3P dehydrogenases, and glycerol kinase suggest derived features of archaeal membranes having G1P polar lipids. Archaea 2016, (2016).

35. Villanueva, L., Schouten, S. & Damsté, J. S. S. Phylogenomic analysis of lipid biosynthetic genes of Archaea shed light on the ‘lipid divide’. Environ. Microbiol. 19, 54–69 (2017).

36. Li, M. et al. Genomic and transcriptomic evidence for scavenging of diverse organic compounds by widespread deep-sea archaea. Nat. Commun. 6, 8933 (2015).

37. Orsi, W. D., Edgcomb, V. P., Christman, G. D. & Biddle, J. F. Gene expression in the deep biosphere. Nature 499, 205 (2013).

38. Lloyd, K. G. et al. Phylogenetically novel uncultured microbial cells dominate Earth microbiomes. MSystems 3, e00055–18 (2018).

39. Bergkessel, M., Basta, D. W. & Newman, D. K. The physiology of growth arrest: uniting molecular and environmental microbiology. Nat. Rev. Microbiol. 14, 549 (2016).

40. Burdige, D. J. Preservation of Organic Matter in Marine Sediments: Controls, Mechanisms, and an Imbalance in Sediment Organic Carbon Budgets? Chem. Rev. 107, 467–485 (2007).

41. Kristensen, E., Bouillon, S., Dittmar, T. & Marchand, C. Organic carbon dynamics in mangrove ecosystems: a review. Auqat. Bot. 89, 201–219 (2008).

42. Sørensen, J., Jørgensen, B. B. & Revsbech, N. P. A comparison of oxygen, nitrate, and sulfate respiration in coastal marine sediments. Microb. Ecol. 5, 105–115 (1979).

43. Zhang, Y. & Zhai, W.-D. Shallow-ocean methane leakage and degassing to the atmosphere: triggered by offshore oil-gas and methane hydrate explorations. Front. Mar. Sci. 2, (2015).

44. Jørgensen, S. L. et al. Quantitative and phylogenetic study of the Deep Sea Archaeal Group in sediments of the Arctic mid-ocean spreading ridge. Front. Microbiol. 4, 299 (2013).

45. Pan, J. et al. Vertical Distribution of Bathyarchaeotal Communities in Mangrove Wetlands Suggests Distinct Niche Preference of Bathyarchaeota Subgroup 6. Microb. Ecol. 77, 417–428 (2019).

46. Zhou, Z. et al. Genomic and transcriptomic insights into the ecology and metabolism of benthic archaeal cosmopolitan, Thermoprofundales (MBG-D archaea). ISME J., 1 (2018).

47. Biddle, J. F. et al. Heterotrophic Archaea dominate sedimentary subsurface ecosystems off Peru. Proc. Natl. Acad. Sci. USA 103, 3846–3851 (2006).

48. Zhang, C. J. et al. Prokaryotic Diversity in Mangrove Sediments across Southeastern China Fundamentally Differs from That in Other Biomes. mSystems 4, e00442–19 (2019).

49. Zhou, Z. et al. Stratified bacterial and archaeal community in mangrove and intertidal wetland mudflats revealed by high throughput 16S rRNA gene sequencing. Front. Microbiol. 8, (2017).

50. Joshi, N. & Fass, J. Sickle: A sliding-window, adaptive, quality-based trimming tool for FastQ files (Version 1.33)[Software]. (2011).

51. Kopylova, E., Noé, L. & Touzet, H. SortMeRNA: fast and accurate filtering of ribosomal RNAs in metatranscriptomic data. Bioinformatics 28, 3211–3217 (2012).

52. Peng, Y., Leung, H. C., Yiu, S. M. & Chin, F. Y. IDBA-UD: a de novo assembler for single-cell and metagenomic sequencing data with highly uneven depth. Bioinformatics 28, 1420–1428 (2012).

53. Bankevich, A. et al. SPAdes: a new genome assembly algorithm and its applications to single-cell sequencing. J. Comput. Biol. 19, 455–477 (2012).

54. Delmont, T. O. & Eren, A. M. Linking pangenomes and metagenomes: the Prochlorococcus metapangenome. PeerJ 6, e4320 (2018).

55. Parks, D. H. et al. CheckM: assessing the quality of microbial genomes recovered from isolates, single cells, and metagenomes. Genome Res. 25, 1043–1055 (2015).

56. Hyatt, D. et al. Prodigal: prokaryotic gene recognition and translation initiation site identification. BMC Bioinformatics 11, 119 (2010).

57. Kanehisa, M., Sato, Y. & Morishima, K. BlastKOALA and GhostKOALA: KEGG tools for functional characterization of genome and metagenome sequences. J. Mol. Biol. 428, 726–731 (2016).

58. Huerta-Cepas, J. et al. Fast genome-wide functional annotation through orthology assignment by eggNOG-mapper. Mol. Biol. Evol. 34, 2115–2122 (2017).

59. Jones, P. et al. InterProScan 5: genome-scale protein function classification. Bioinformatics 30, 1236–1240 (2014).

60. Rawlings, N. D., Barrett, A. J. & Finn, R. Twenty years of the MEROPS database of proteolytic enzymes, their substrates and inhibitors. Nucleic Acids Res. 44, D343–D350 (2015).

61. Bagos, P. et al. Prediction of signal peptides in archaea. Protein Eng. Des. Sel. 22, 27–35 (2008).

62. Yu, N. Y. et al. PSORTb 3.0: improved protein subcellular localization prediction with refined localization subcategories and predictive capabilities for all prokaryotes. Bioinformatics 26, 1608–1615 (2010).

63. Vanwonterghem, I. et al. Methylotrophic methanogenesis discovered in the archaeal phylum Verstraetearchaeota. Nat. Microbiol. 1, 16170 (2016).

64. Spang, A., Caceres, E. F. & Ettema, T. J. Genomic exploration of the diversity, ecology, and evolution of the archaeal domain of life. Science 357, eaaf3883 (2017).

65. Durbin, A. M. & Teske, A. Archaea in organic-lean and organic-rich marine subsurface sediments: an environmental gradient reflected in distinct phylogenetic lineages. Front. Microbiol. 3, 168 (2012).

66. Pruesse, E., Peplies, J. & Glöckner, F. O. SINA: accurate high-throughput multiple sequence alignment of ribosomal RNA genes. Bioinformatics 28, 1823–1829 (2012).

67. Nguyen, L. T., Schmidt, H. A., von Haeseler, A. & Minh, B. Q. IQ-TREE: a fast and effective stochastic algorithm for estimating maximum-likelihood phylogenies. Mol. Biol. Evol. 32, 268–274 (2014).

68. Mistry, J. et al. Challenges in homology search: HMMER3 and convergent evolution of coiled-coil regions. Nucleic Acids Res. 41, e121 (2013).

69. Eddy, S. R. Accelerated Profile HMM Searches. PLoS Comput. Biol. 7, e1002195 (2011).

70. Criscuolo, A. & Gribaldo, S. BMGE (Block Mapping and Gathering with Entropy): a new software for selection of phylogenetic informative regions from multiple sequence alignments. BMC Evol. Biol. 10, 210 (2010).

71. Caspi, R. et al. The MetaCyc Database of metabolic pathways and enzymes and the BioCyc collection of Pathway/Genome Databases. Nucleic Acids Res. 36, D623–D631 (2007).

72. Li, H. & Durbin, R. Fast and accurate short read alignment with Burrows–Wheeler transform. Bioinformatics 25, 1754–1760 (2009).

