## supplementary materials 1 for "Highly diverse Asgard archaea participate in organic matter degradation in coastal sediments"

**This file includes:**

Methods
Asgard archaea 16S rRNA gene dataset construction
Phylogenetic position and distribution of Asgard archaea 16S rRNA gene sequences
Molecular modelling and dynamics simulation
Evolutionary analysis

Results
Asgard archaea are diverse
Alkane metabolism in Helarchaeota
Genomic and transcriptomic evidence for carbon fixation in Asgard archaea

Fig. S1. Amino acid identity (AAI) of Asgard archaea MAGs.
Fig. S2. Pan-genome analysis of protein clusters within all Asgard MAGs using the Anvi'o software.
Fig. S3. Non-metric multidimensional scaling (NMDS) plot based on Bray–Curtis dissimilarities for KEGG annotations in all available Asgard MAGs.
Fig. S4. Maximum likelihood phylogenetic analyses of Topoisomerase IB, ribophorin I, and DNA-directed RNA polymerase A.
Fig. S5. Key potential metabolic pathways of Asgard archaea.
Fig. S6. Phylogenetic position of the Gerdarchaeota cytochrome *c* oxidases.
Fig. S7. Phylogenetic position of the Gerdarchaeota rhodopsins.
Fig. S8. Abundance of (A) peptidases and (B) carbohydrate-active enzymes in Asgard MAGs.
Fig. S9. Phylogenetic position of the Gerdarchaeota [NiFe]-hydrogenases.
Fig. S10. Phylogenetic maximum likelihood tree of RuBisCO amino acid sequences (large subunit).
Fig. S11. Diversity and biotope of Asgard archaea.
Fig. S12. Protein tree based on *mcrA* gene sequences as identified in the scaffolds.
Fig. S13. Molecular modelling and dynamics of MCR complex.
Fig. S14. Phylogenetic position and evolution of Asgard archaea *mcrA* genes.
Fig. S15. Protein trees of Asgard archaea (A) McrB and (B) McrG.
Fig. S16. Phylogenetic position of the Gerdarchaeota *nifH*.

References (73–98)

### Methods

#### Asgard archaea 16S rRNA gene dataset construction

Archaeal 16S rRNA gene sequences were retrieved from the GenBank NCBI nucleotide database (September 2017) and SILVA SSU r132 database[73]. E-utilities[74] was applied to search and retrieve the archaeal 16S rRNA gene sequences from the NCBI nucleotide database using the ESearch function with the following string: "16S AND 800:2000[Sequence Length] AND archaea[organism] AND rrna[Feature Key] AND isolation_source[All fields] NOT genome NOT chromosome NOT plasmid". EFetch function was then used to retrieve the sequences and the corresponding GenBank-formatted flat file, which contained environmental information (e.g., location and isolation source). In the subsequent steps, custom scripts were designed to combine the two datasets and to remove low-quality (i.e., containing 'N' or shorter than 800 bp) and duplicate sequences, resulting in 100,786 archaeal sequences. To retrieve Asgard 16S rRNA gene sequences, the above obtained sequences were BLAST-searched against genome-based 16S rRNA gene sequences (≥800 bp) using BLASTn with a cutoff E-value ≤1e-5, sequence identity ≥75%, and coverage ≥50%. This resulted in a set of 9765 potential Asgard sequences from the public databases. OTUs were assigned using the QIIME UCLUST[75] wrapper, with a threshold of 95% nucleotide sequence identity, and the cluster centroid for each OTU was chosen as the representative OTU sequence. A set of OTU threshold (e.g., 90%, 95% and 97%) was verified to obtain the optimum value for phylogenetic tree building and data analysis.

In addition to public databases, archaeal 16S rRNA gene sequences were also retrieved from a recent study[76]. All expressed 16S rRNA gene sequences in the reference paper were BLASTn-searched against a custom database containing 16S rRNA gene sequences retrieved from Asgard MAGs and potential Asgard OTUs obtained as described above, with a cutoff E-value ≤1e-5, sequence identity ≥50%, and coverage ≥80%. Finally, 5588 potential Asgard archaeal sequences were obtained, including eight newly proposed clusters DAS1–8 (79 sequences)[76]. OTU re-formation of these sequences (15,353 sequences, from both databases and the reference paper) was performed using the QIIME UCLUST[75] wrapper, with a threshold of 95% pair-wise nucleotide sequence identity, resulting in 1836 OTUs.

#### Phylogenetic position and distribution of Asgard archaea 16S rRNA gene sequences

SINA-aligned[66] archaeal representative OTU sequences obtained in the previous step (1836 OTUs) were pre-filtered using a backbone tree, which was constructed based on updated 16S rRNA archaeal gene datasets[64, 65], using the ARB software (version 5.5) with the "Parsimony (Quick Add Marked)" tool[77]. This resulted in 456 Asgard OTUs. Candidate representative OTU sequences (456 OTUs), genome-based 16S rRNA gene sequences ≥800 bp (17 sequences), and reference sequences were used for phylogenetic tree construction. Maximum-likelihood tree was inferred with IQ-TREE (version 1.6.1)[67] using the GTR+I+G4 mixture model (recommended by the "TESTONLY" model) and ultrafast (-bb 1000). Asgard clade designations were made when either a group was defined in previous publications, or when groups with >20 sequences and intragroup similarity >83%[78] were monophyletic. Attributes [i.e., expressed rRNA gene, biotope, temperature, and salinity] for each representative OTU were extracted and visualized using iTOL software[79]. Calculation of percent identity of new Asgard clades was based on 179 sequences with a long fragment of Asgard archaeal 16S rRNA gene. Fragments of 16S rRNA gene from position of *E. coli* 243 to 1414 (~1170 bp) were used for calculating the identity.

The corresponding environmental information (i.e., location and biotopes) for the 16S rRNA gene sequences of Asgard archaea was extracted from the GenBank-formatted flat file using custom scripts. This resulted in 172 libraries with latitude and longitude of sampling sites (table S9). Information on the sample locations was plotted using the mapdata and ggplot2 packages in R software.

**Molecular modelling and dynamics simulation**

Lokiarchaeotal MCR amino acid sequences were blasted against the protein data bank (PDB)[80] to obtain high-similarity sequences. The geometry of these sequences was then predicted using MODELLER[81], and those with high DOPE score were kept for analysis. Protein–protein interaction of the MCR complex was predicted using ZDOCK (http://zdock.umassmed.edu/), and the protein–ligand docking was executed by the AutoDock4 (version 4.2.6)[82] with the Lamarckian genetic algorithm[83].

Molecular dynamics (MD) simulations of MCR complexes were performed using GROMACS (version 5.1.1)[84] based on AMBER99SB-ILDN force field[85] and TIP3P water box[86]. The cutoff of non-bonded interactions involving van der Waals and electrostatics was set 10 Å. Long range electrostatic interactions were treated using the Partice-Mesh-Ewald (PME) algorithm[87]. Energy minimization was carried out to obtain an initial structure with force ≤1000 kJ·mol-1·nm-1. MD simulations were initially carried out using the NVT ensemble for 100 ps (300 K), followed by simulation with NPT ensemble for 100 ps (300 K and 1 bar). Then, 50 ns MD simulations were performed using NPT ensemble (300 K and 1 bar) with the time steps of 2 fs to get the equilibrium trajectories. The snapshot was saved at an interval of 10 ps for subsequent analysis. Snapshots were saved every 10 ps for subsequent analysis.

**Evolutionary analysis**

All genomes containing MCR complex were downloaded from NCBI (https://www.ncbi.nlm.nih.gov/) databases. CheckM was used to check the genome quality. Genomes with completeness < 70% were removed for further analysis. We selected 117 genomes for final analysis. The phylogenetic tree of concatenated 122 archaeal marker genes were constructed using IQ-TREE (version 1.6.3) with "-m MFP -mset LG,WAG -mrate E,I,G,I+G -mfreq FU -bb 1000" flags, the genome of a DPANN archaea (Diapherotrites_AR10) was used as outgroup. Putative HGTs were inferred using HGTector[88]. A standard database (version updated on 2017-6-30) were used for homologues searching by using MMseqs2[89]. Quality cutoffs for valid hits were e-value ≤1e-20, sequence identity ≥30%, and coverage of query sequence ≥50%.

### Results

**Asgard archaea are diverse**

To get an overview of the diversity of Asgard archaea, 16S rRNA gene sequences (> 600 bp) from all Asgard metagenome-assembled genomes (MAGs) and public databases (SILVA SSU 132 and NCBI databases), including a recent study of 16S rRNA gene sequences[76], were systematically analysed. After sequence filtering, >400 Asgard archaeal operational taxonomic units (OTUs, 95% cutoff, 10,448 sequences) were obtained. Based on the phylogenetic analysis, five new Asgard lineages at phylum or class level (68-85% sequence similarity to other known Asgard phyla, tables S3 and S12) were identified. Intriguingly, most sequences (99.7%) of these new groups were from expressed 16S rRNA genes[76], indicating that Asgard archaea are more diverse than previously proposed[3]. Besides, with a lower minimum intragroup similarity (71%, table S12) than the recommended phylum-level threshold (75%)[90], the lineage of

Heimdallarchaeota was divided into two subgroups: Heimdallarchaeota-AAG and Heimdallarchaeota-MHVG, following previous suggestions[3, 91].

**Alkane metabolism in Helarchaeota**

In agreement with a previous report[4], the genes encoding Helarchaeotal methyl-coenzyme M reductase (MCR) found in this study clustered together within the branch of the butane-oxidizing Syntrophoarchaeum[92] and ethane-oxidizing Argoarchaeum[93] (figs. S14A and S15). The expressed Helarchaeota-like *mcrA* genes in unbinned scaffolds (e.g., SZ_4_scaffold_203331_2, fig. S12) highlight the involvement of Helarchaeota in activities of alkane oxidation in coastal sediments. Molecular modelling and dynamics studies supported this relatedness by showing that the MCR complex of Helarchaeota for butane binding was similar to that of *Ca*. Syntrophoarchaeum according to the root mean square deviation (RMSD) values (~5 Å, fig. S13). Thus, Helarchaeota might be able to oxidize short-chain alkanes, in which ethane and butane are preferentially used (fig. S13). In addition, based on evolutionary analysis, a high percentage (> 10%) of *mcrA* genes in the Helarchaeota MAGs was most likely transferred horizontally from archaea, and most of these genes originated from methanogenic hosts/donors, e.g., Thermococci, Methanomicrobia, and Methanobacteria (fig. S14B).

**Metabolic potentials of Gerdarchaeota**

Gerdarchaeota MAGs contain genes encoding *nifH* for potential nitrogen metabolism but with no genes for *nifD* and *nifK* (Fig. 2). Phylogenetic analysis indicates that the annotated Gerdarchaeotal *nifH* clusters with homologs found in other Asgard archaea and belongs to the type IV category together with methanogens, although they are distinct to the known type IV *nifH* (fig. S16), which is required for the biosynthesis of cofactor F430[94, 95]. Thus, Asgard archaea might rather not perform nitrogen fixation.

**Genomic and transcriptomic evidence for carbon fixation in Asgard archaea**

Differ to Heimdallarchaeota-AAG and Heimdallarchaeota-MHVG, the complete gene sets for the Wood-Ljungdahl (WL) pathway in Lokiarchaeota and Thorarchaeota MAGs implies that they may use both tetrahydrofolate (THF) and tetrahydromethanopterin (THMPT) as $C_1$ carriers. Intriguingly, transcripts for the THMPT-dependent WL pathway were predominant, especially those for incorporating $CO_2$ into formyl-methanofuran and $CO_2$ reduction to acetyl-CoA by methyl-THMPT, showing that the THMPT-dependent WL pathway might be preferred in these archaea. As for Helarchaeota, they may use THMPT as C1 carrier for carbon fixation as revealed by the transcripts. Furthermore, generation of pyruvate from acetyl-CoA and $CO_2$ via an anaplerotic pathway elevated the contribution of inorganic carbon to biomass for all phyla[33]. Additionally, genomic and transcriptomic evidence suggest that Lokiarchaeota, Heimdallarchaeota-AAG and Heimdallarchaeota-MHVG have carbon fixation potential via the incomplete rTCA cycle. The equipped gene sets for $CO_2$ assimilation ensure Asgard arcgaea to survive in the oligotrophic biosphere (e.g., deep subsurface marine sediment) where the energy supply is extraordinarily low but inorganic carbon is sufficient[96, 97]. The versatile abilities to perform inorganic carbon assimilation and organics degradation (aerobically and anaerobically) may facilitate these species response and adapt to environmental changes immediately.

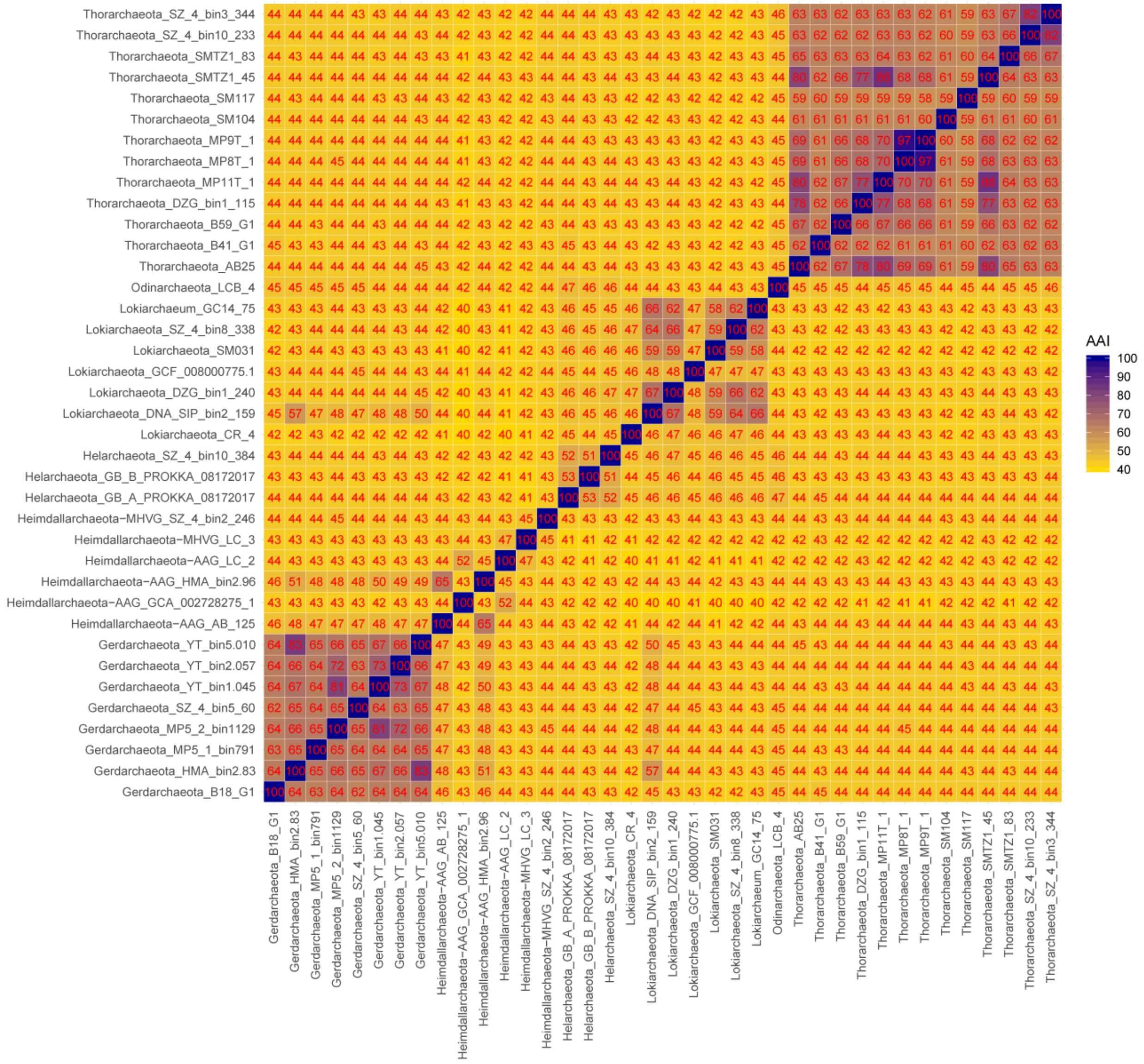

**Fig. S1. Amino acid identity (AAI) of Asgard archaea MAGs.** AAI was calculated using CompareM (https://github.com/dparks1134/CompareM).

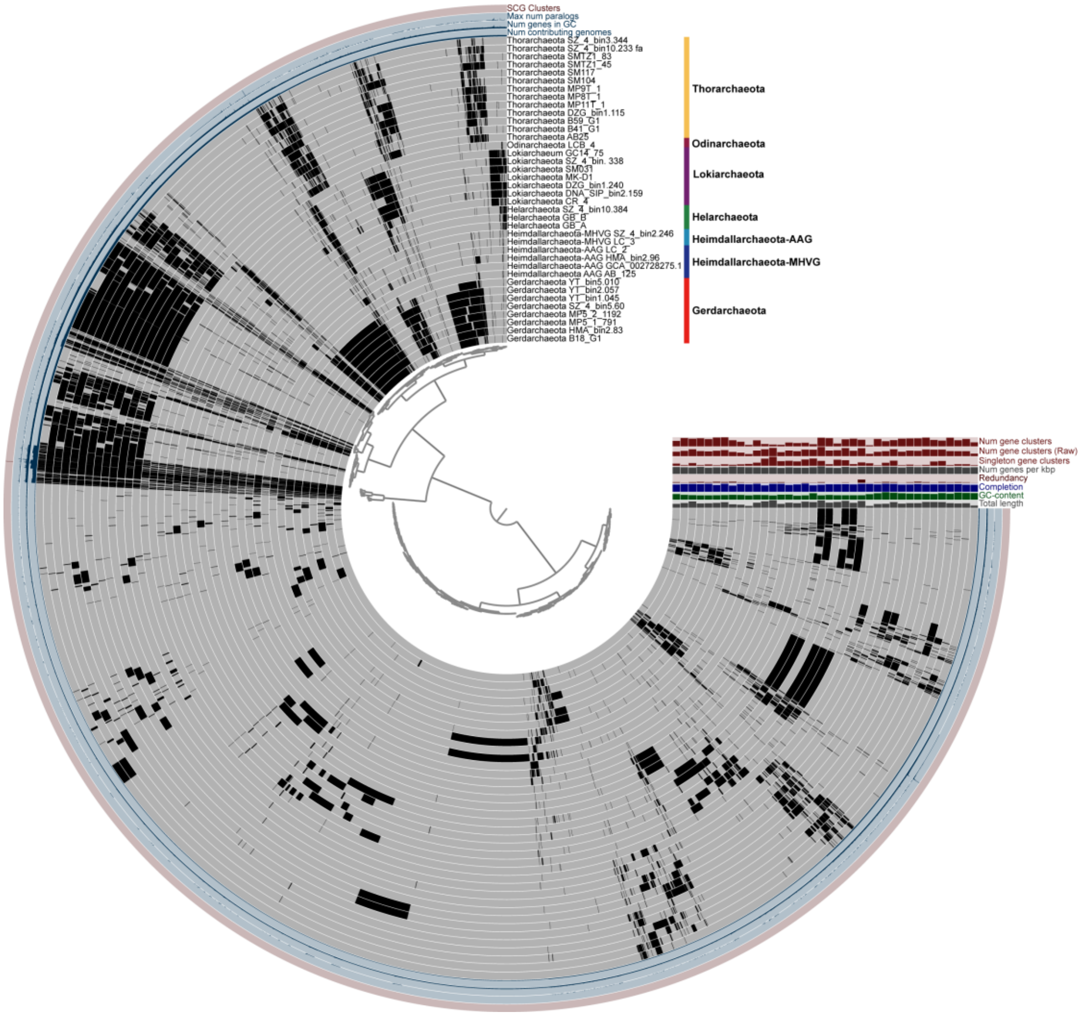


**Fig. S2. Pan-genome analysis of protein clusters within all Asgard MAGs using the Anvi'o software**. The inner tree was clustered using the function "Gene cluster presence absence". Genes for comparison were selected using the option "--min-occurrence 3". Black and gray color represents presence and absence, respectively. The brown bar denotes the cluster of COGs for the single conserved genes (SCGs) that were shared by all the genomes.

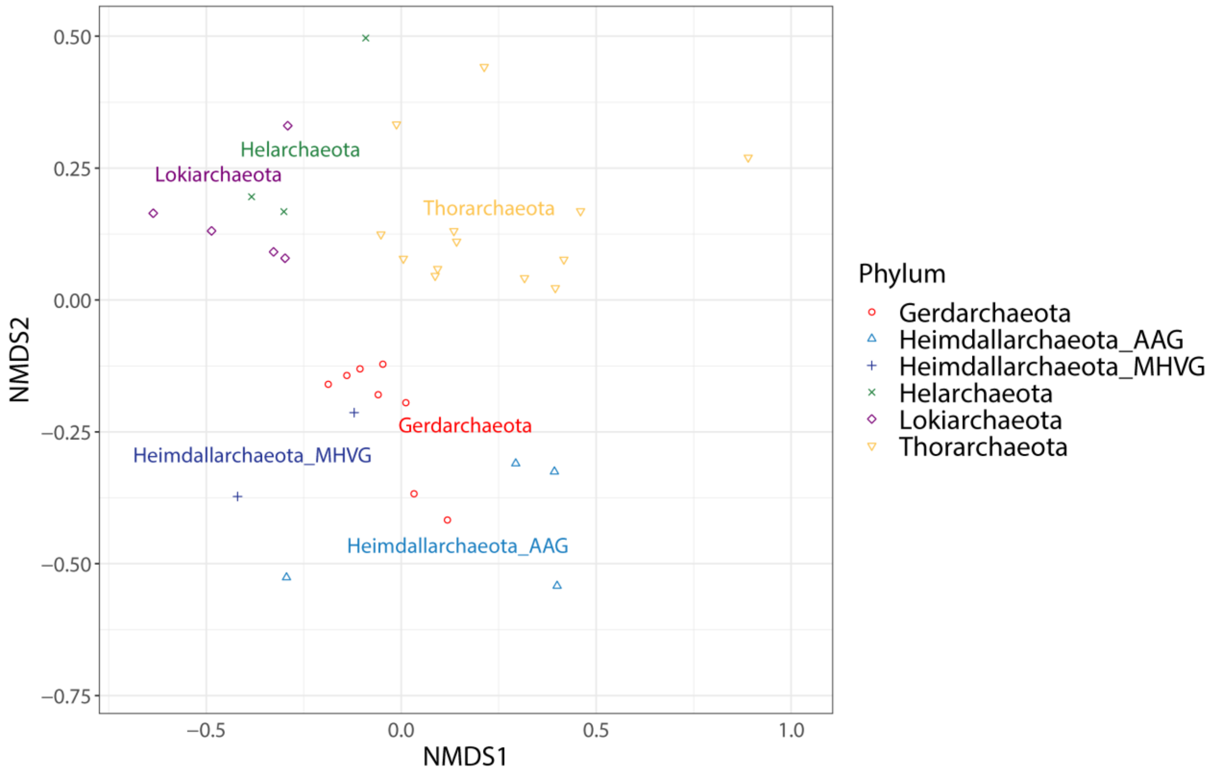


**Fig. S3. Non-metric multidimensional scaling (NMDS) plot based on Bray–Curtis dissimilarities for KEGG annotations in all available Asgard MAGs.** Odinarchaeota was excluded since it was only represented by one MAG.

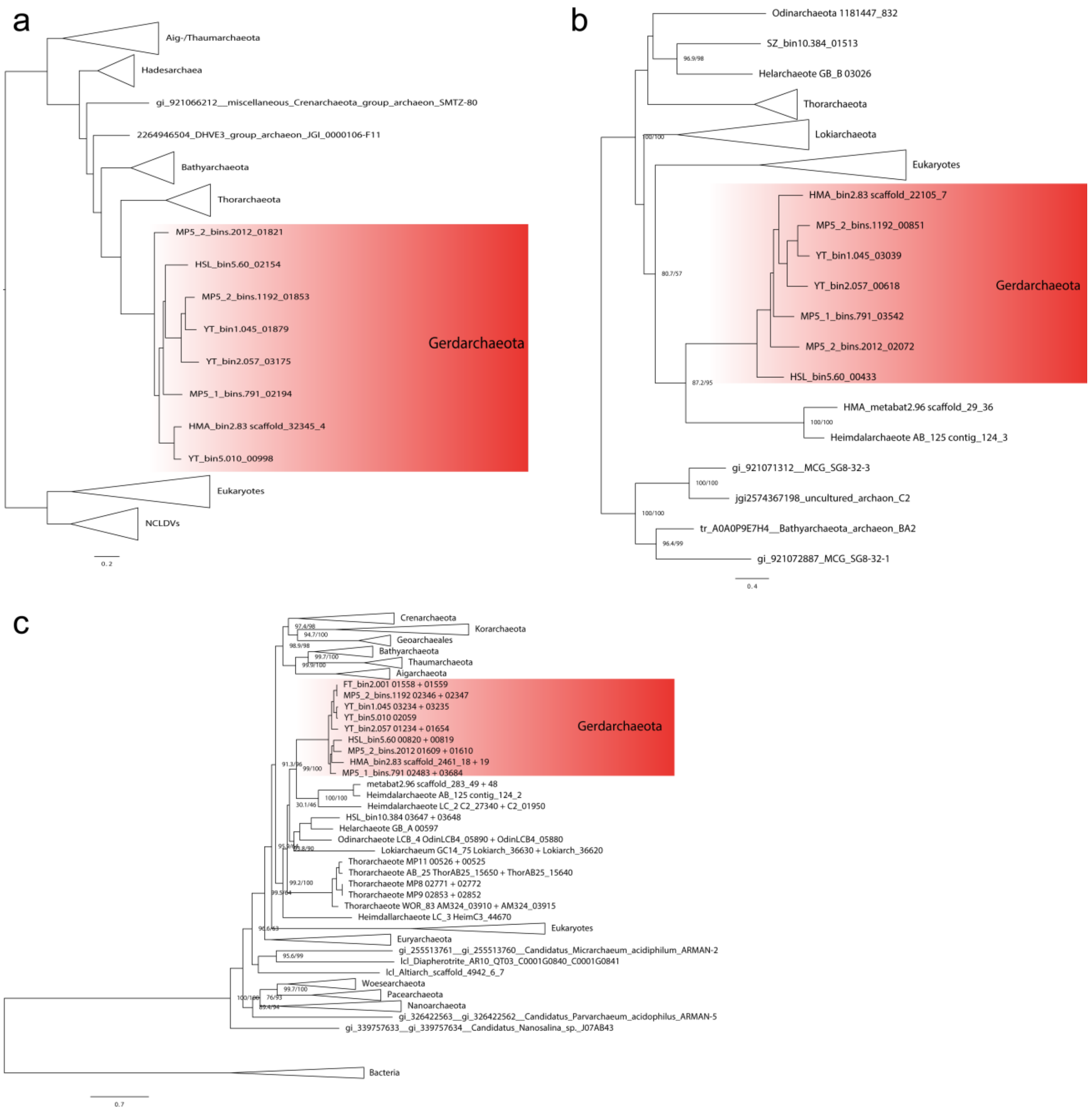


**Fig. S4. Maximum likelihood phylogenetic analyses of Topoisomerase IB, ribophorin I, and DNA-directed RNA polymerase A.** (**A**), Unrooted Topoisomerase IB phylogeny inferred from an alignment consisting of 501 amino acid positions with LG4X model. (**B**), ribophorin I phylogeny inferred from an alignment consisting of 547 amino acid positions with LG+F+R4 model. (c) DNA-directed RNA polymerase A phylogeny inferred from an alignment consisting of 848 amino acid positions with LG+C20+G model. Bold characters indicate the lineages with no fused genes.

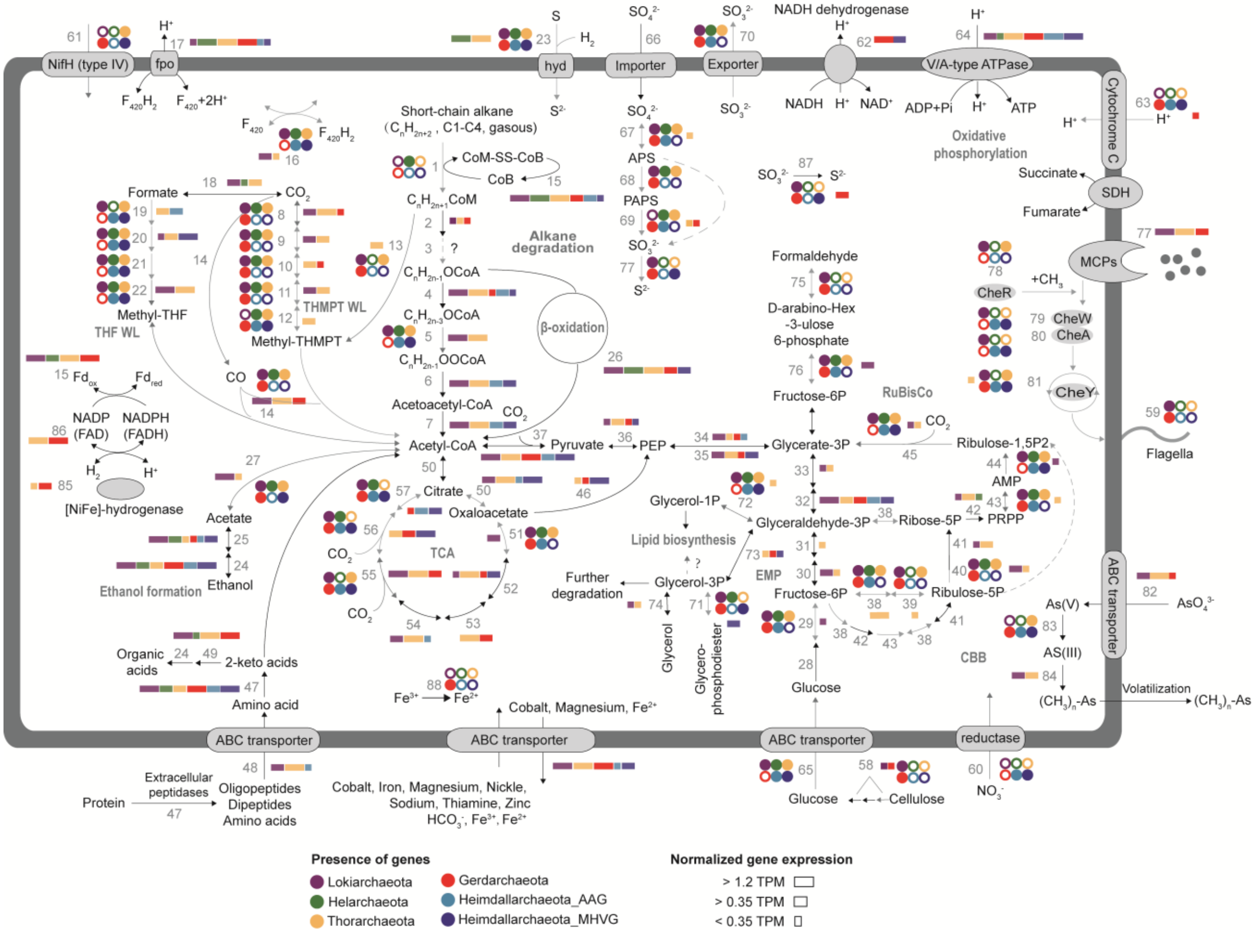


**Fig. S5. Key potential metabolic pathways of Asgard archaea.** The whole pathway was reconstructed based on all available Asgard MAGs (table S2). Circles with different colours represent different phyla or MAGs, with filled circles indicate presence of genes. For comparison purposes, we used arrows of different types and colours. Black arrows indicate genes found in all Asgard MAGs, and grey arrows represent genes that are present in a subset of MAGs. Dashed grey arrows show pathways that are missing from all MAGs. Detailed metabolic information and transcript abundance for surface and subsurface sediments are available in tables S6 and S7. The relative abundance of the transcripts (Transcripts Per Million reads, TPM) for each gene is marked with proportionally sized rectangles and represented at the phylum level. Odinarchaeota were excluded because no Odinarchaeota MAG was identified in the present study.

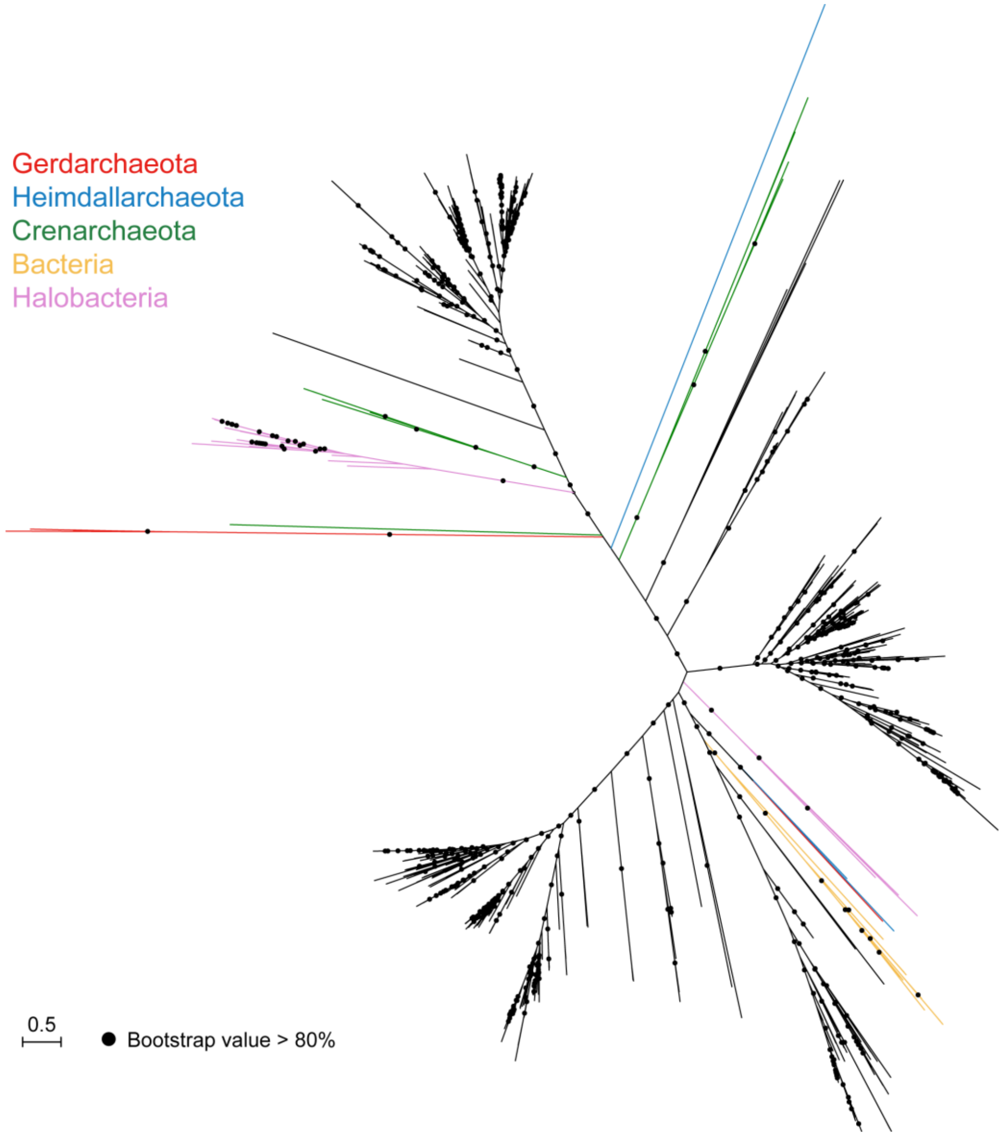


**Fig. S6. Phylogenetic position of the Gerdarchaeota cytochrome *c* oxidases.** The neighbour lineages of Gerdarchaeota cytochrome *c* oxidases are marked with different colours. The unrooted maximum-likelihood tree was obtained using IQ-TREE software with the mixture mode 'WAG+F+G4'.

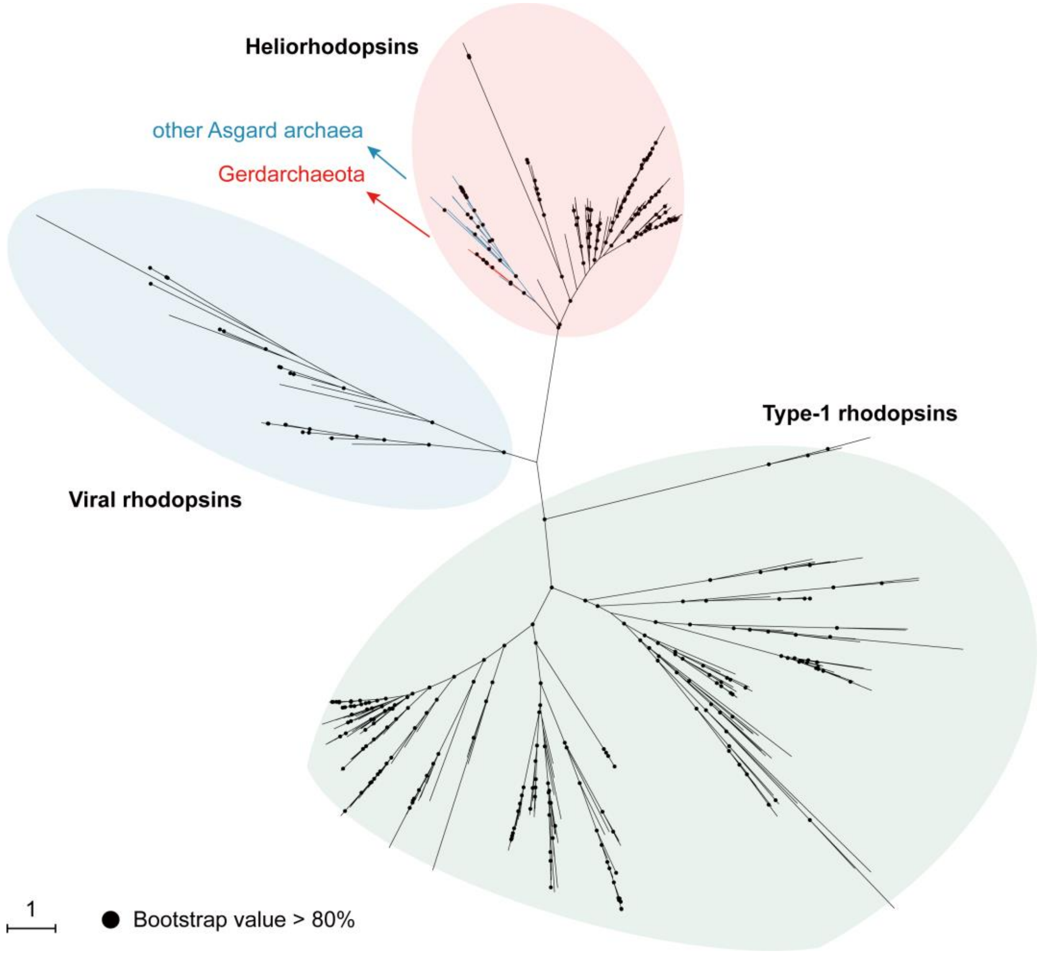


**Fig. S7. Phylogenetic position of the Gerdarchaeota rhodopsins.** The unrooted maximum-likelihood tree was obtained using IQ-TREE software with the mixture mode 'VT+F+R9'.

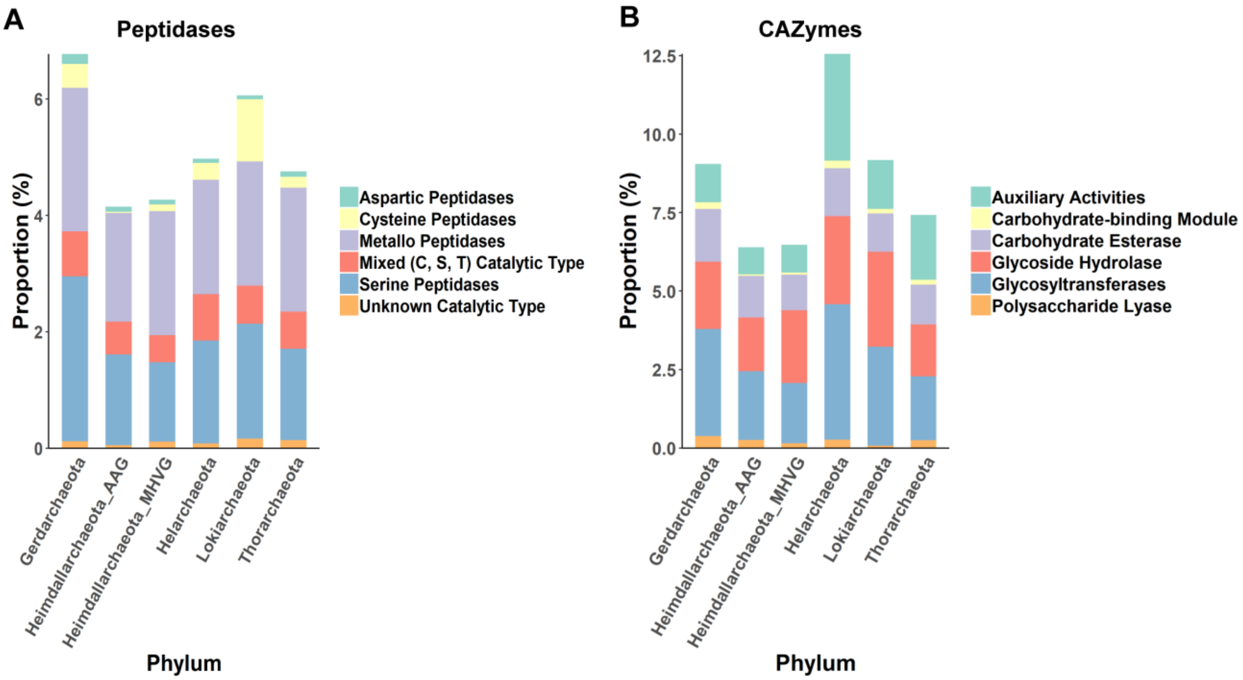


**Fig. S8. Abundance of (A) peptidases and (B) carbohydrate-active enzymes in Asgard MAGs.** Carbohydrate-active enzymes (CAZymes) and peptidases were annotated using the dbCAN webserver and MEROPs database, respectively. The e-value cutoff is 1e-5 for both cases. Proportions were calculated by normalizing the MAGs number and the average protein numbers in each phylum.

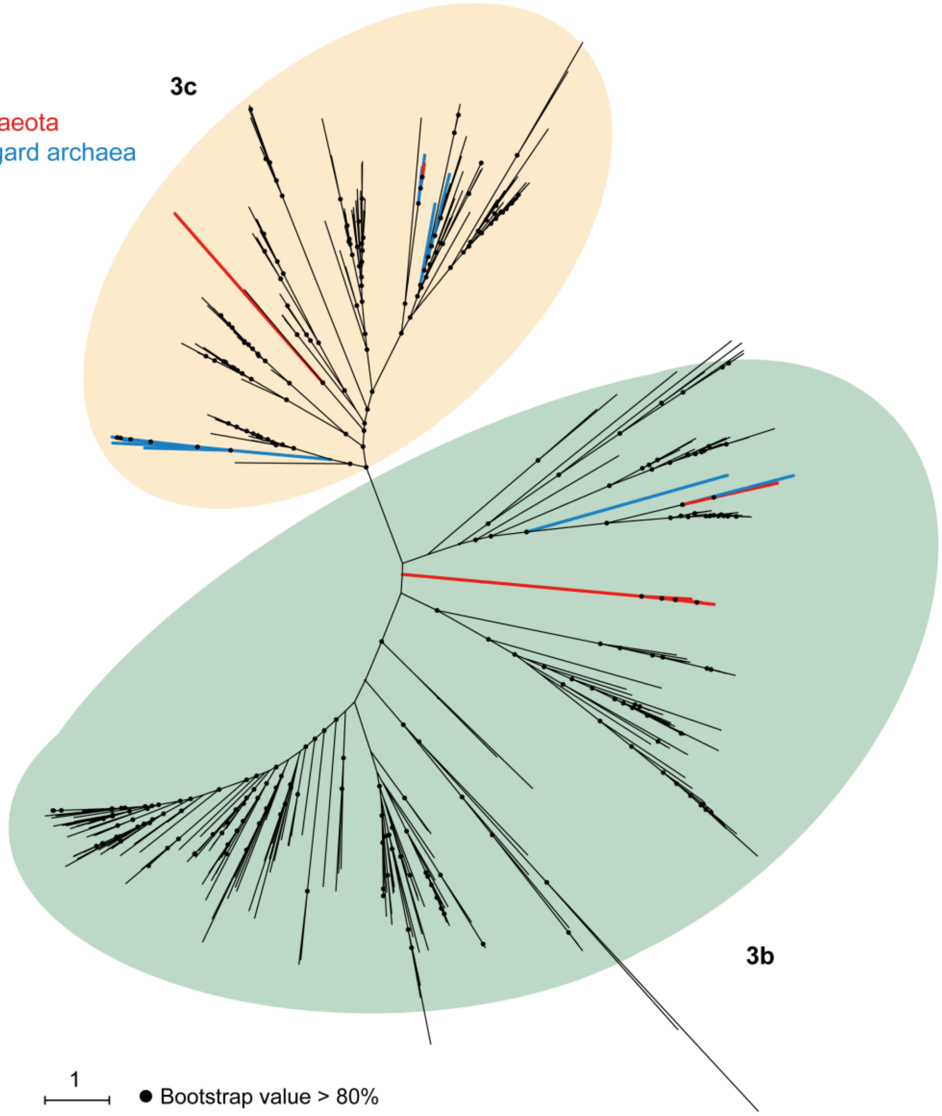


**Fig. S9. Phylogenetic position of the Gerdarchaeota [NiFe]-hydrogenases.** The unrooted maximum-likelihood tree was obtained using IQ-TREE software with the mixture mode 'LG+G4'. Reference sequences for groups 3b and 3c [NiFe]-hydrogenases were obtained based on a reference database HydDB[98].

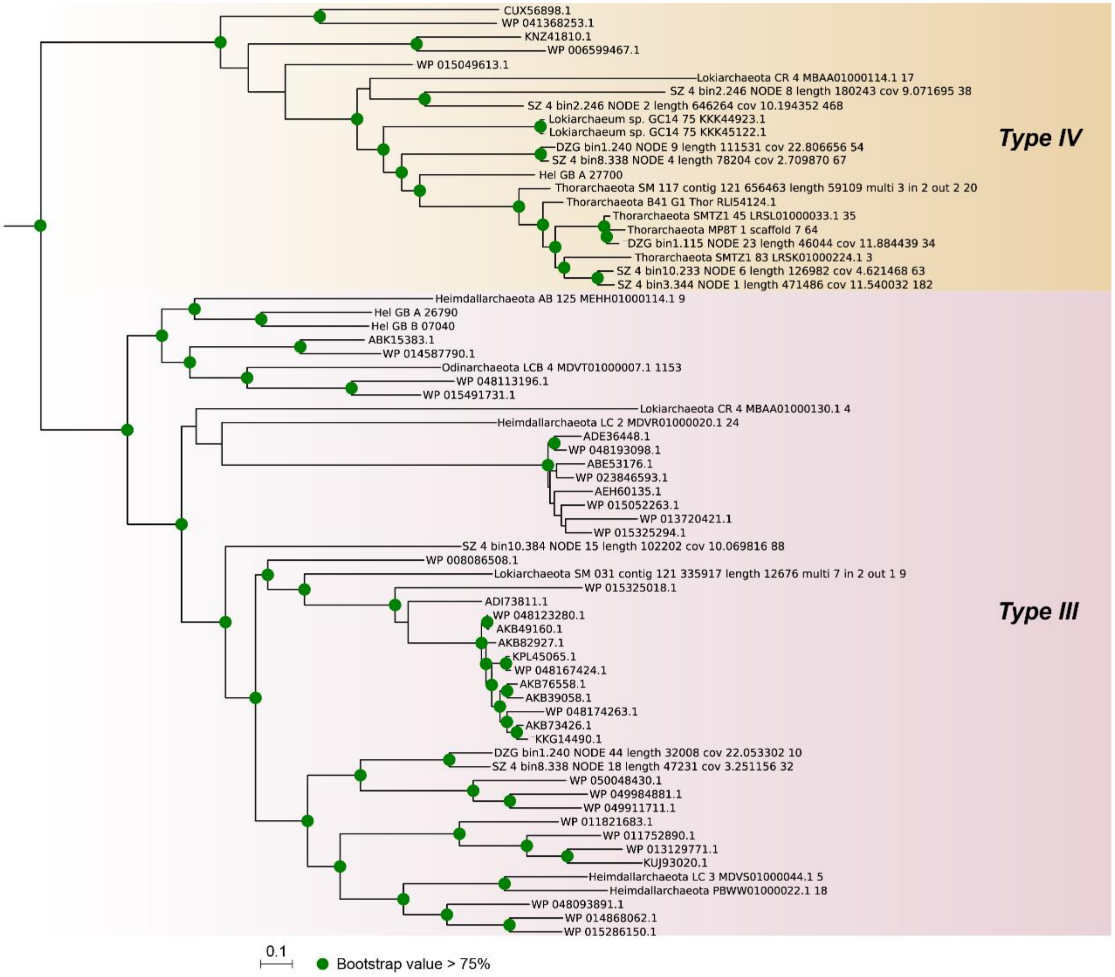


**Fig. S10. Phylogenetic maximum likelihood tree of RuBisCO amino acid sequences (large subunit).** The tree was build using IQ-TREE with model LG+I+G4 and parameter “-bb 1000”.

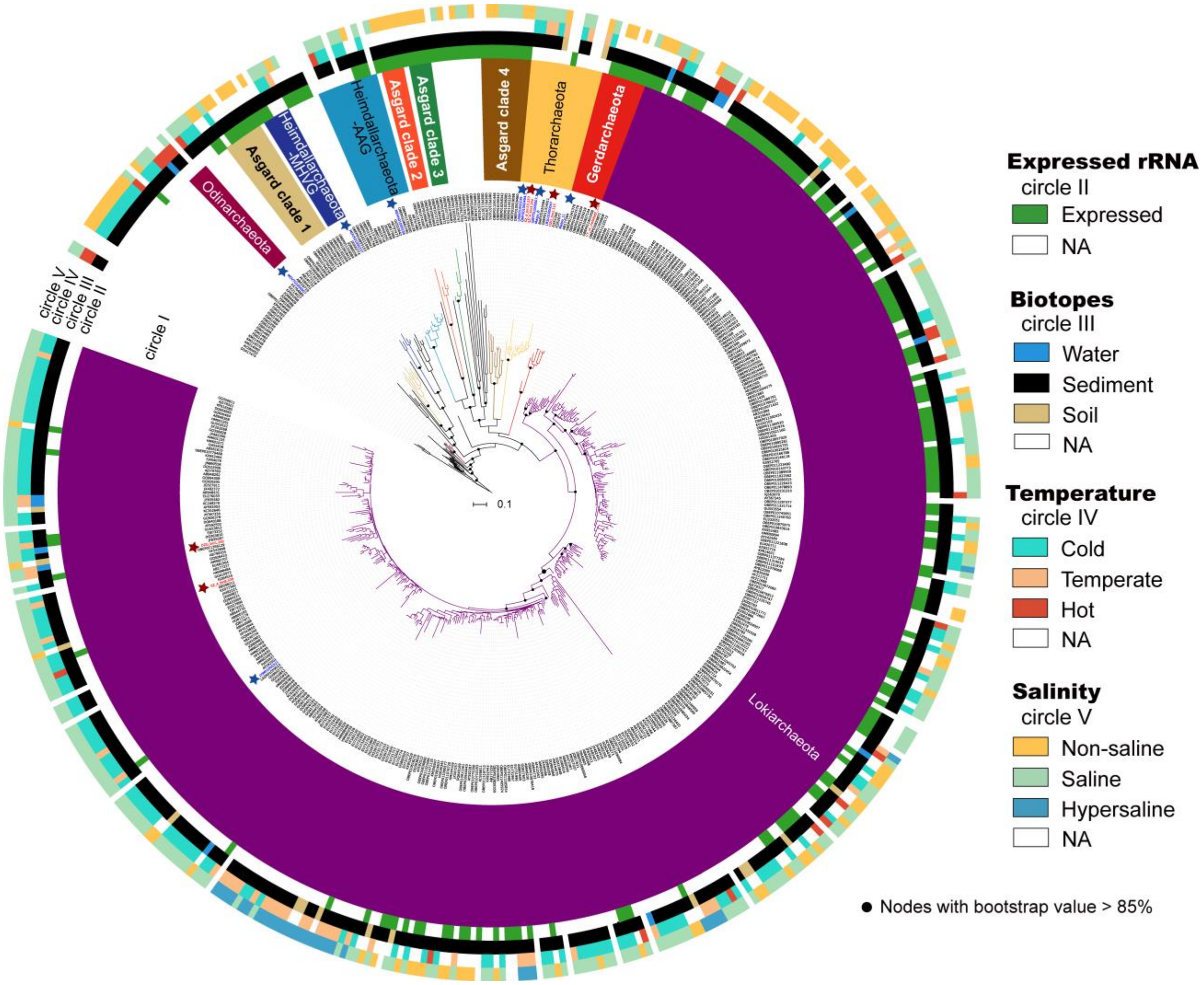


**Fig. S11. Diversity and biotope of Asgard archaea.** Maximum likelihood tree of Asgard archaea constructed using genome-based and publicly available 16S rRNA gene sequences clustered at 95% sequence identity. Groups were designated based on sequences fulfilling the division criteria (see Methods). The colour circles, from the inside to the outside, represent the expressed 16S rRNA gene sequences, habitat, temperature, and salinity, accordingly. Red stars represent 16S rRNA gene sequences from newly discovered Asgard MAGs, and blue stars represent sequences from reference Asgard MAGs. The tree was rooted with Crenarchaeota. Key nodes with bootstrap support values >70% are shown.

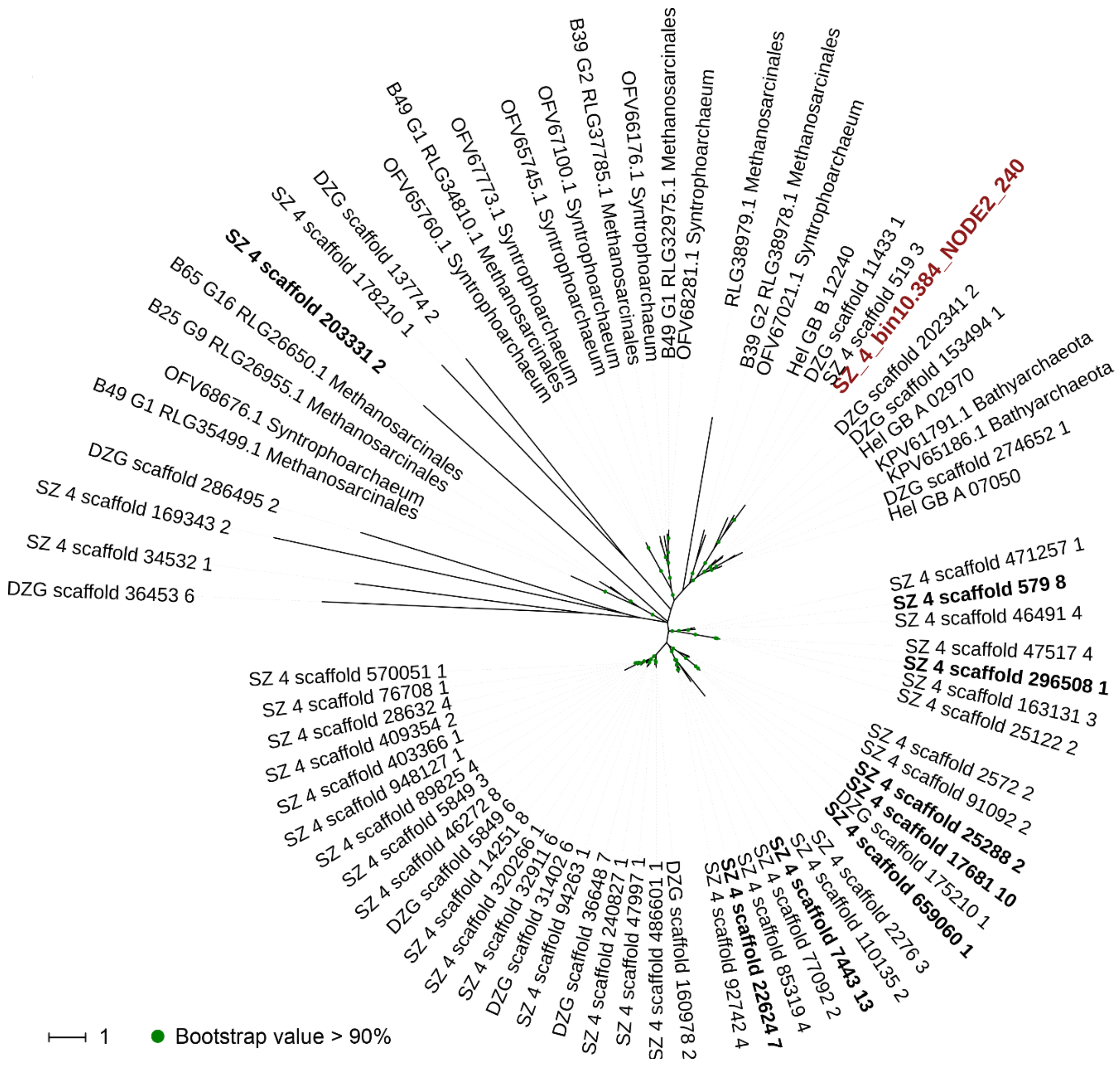


**Fig. S12. Protein tree based on *mcrA* gene sequences as identified in the scaffolds.** The protein sequences were obtained through BLASTP against novel McrA protein sequences of Asgard with E-value cutoff ≤1e-5. Amino acids > 400 aa were kept for tree building. The tree was rooted with Euryarchaeotal and Verstraetearchaeotal McrA protein sequences. McrA sequences deduced from transcripts are marked in bold.

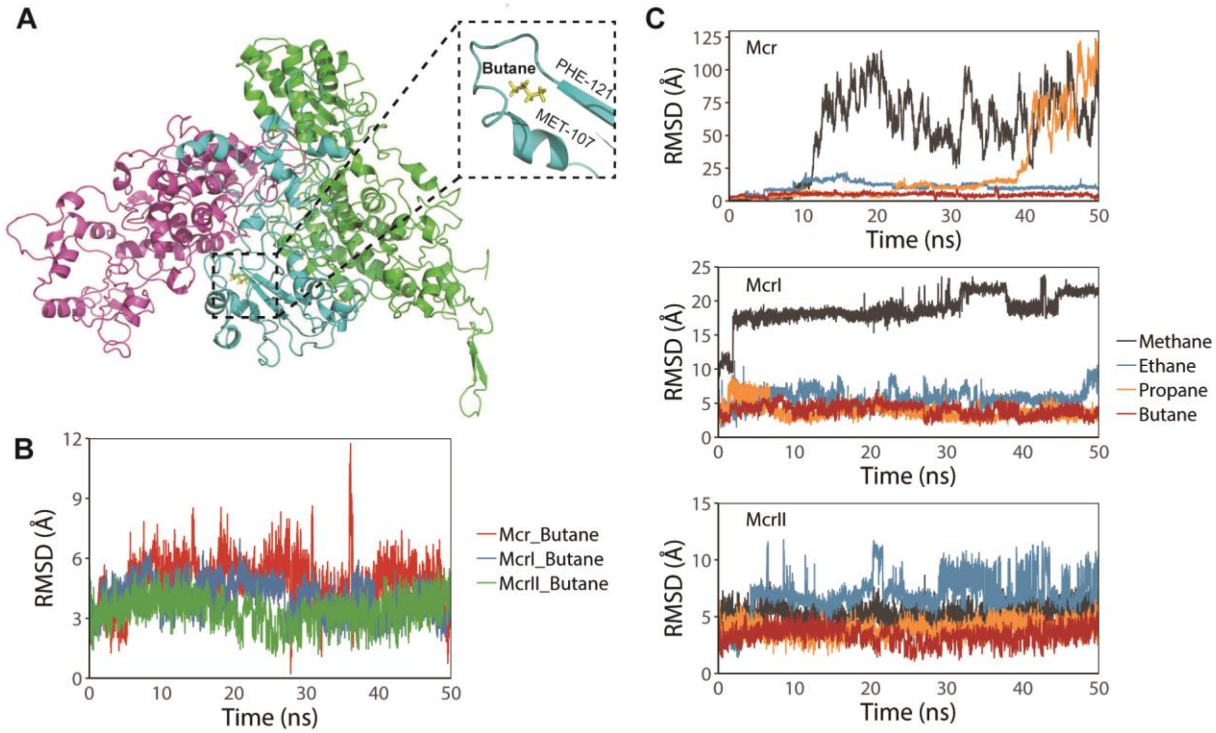


**Fig. S13. Molecular modelling and dynamics of MCR complex.** (**A**), Equilibrium structure of the Lokiarchaeotal MCR complex and the docking model of butane (yellow). McrA, McrB and McrG are marked in green, magenta, and cyan, respectively. Magnified structure of butane binding with the pocket of McrG shown as inset. The main residues interacting with butane are from MET107 to PHE121 of McrG. (**B**), The RMSD of butane in Mcr (red), McrI (blue) and McrII (green) complexes as a function of time. The RMSD of butane in the three complexes are of the same magnitude (~5Å). (**C**), The RMSD of methane (black), ethane (blue), propane (yellow) and methane (red) in Mcr (top), McrI (middle) and McrII (bottom) complexes as a function of time. The RMSD of butane in all the three complexes are of smaller magnitude than the ones of methane, which implies that the binding affinities of butane to all the three Mcr complexes are higher than the ones of methane. Here, Mcr represents the Lokiarchaeotal MCR complex, while McrI and McrII represent the complexes from the *Ca*. Syntrophoarchaeum butanivorans. The RMSD is estimated as the distance deviation of methane/butane to the center of mass of each Mcr complex.

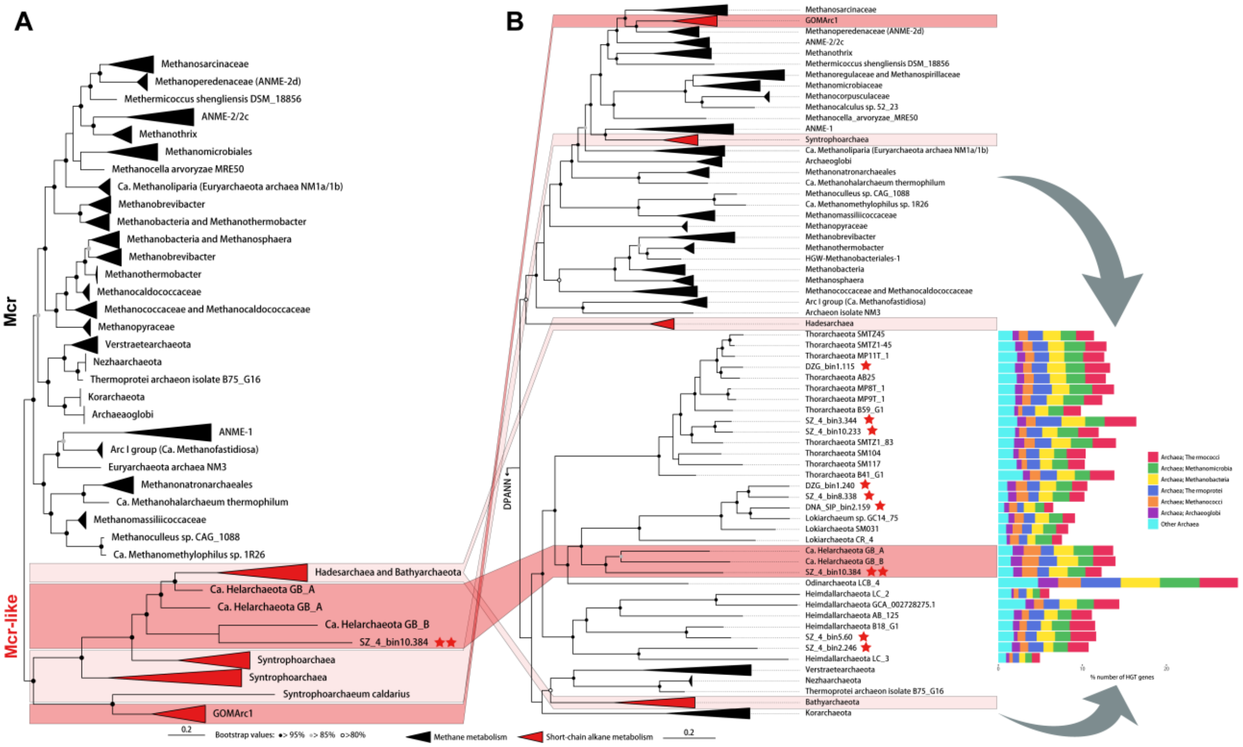


**Fig. S14. Phylogenetic position and evolution of Asgard archaea *mcrA* genes.** (**A**) Maximum likelihood phylogenetic tree of the *mcrA* genes. (**B)** Ancestral genome content reconstruction with the concatenated 122 archaeal marker gene tree inferred from IQ-TREE. Bar plot of the percentage of genes horizontally transferred from archaea to their genomes. The genes and MAGs obtained in this article are marked with red stars, and those with the MCR complex are highlighted with two red stars. The two grey arrows indicate genes that underwent extensive horizontal transfer from archaea.

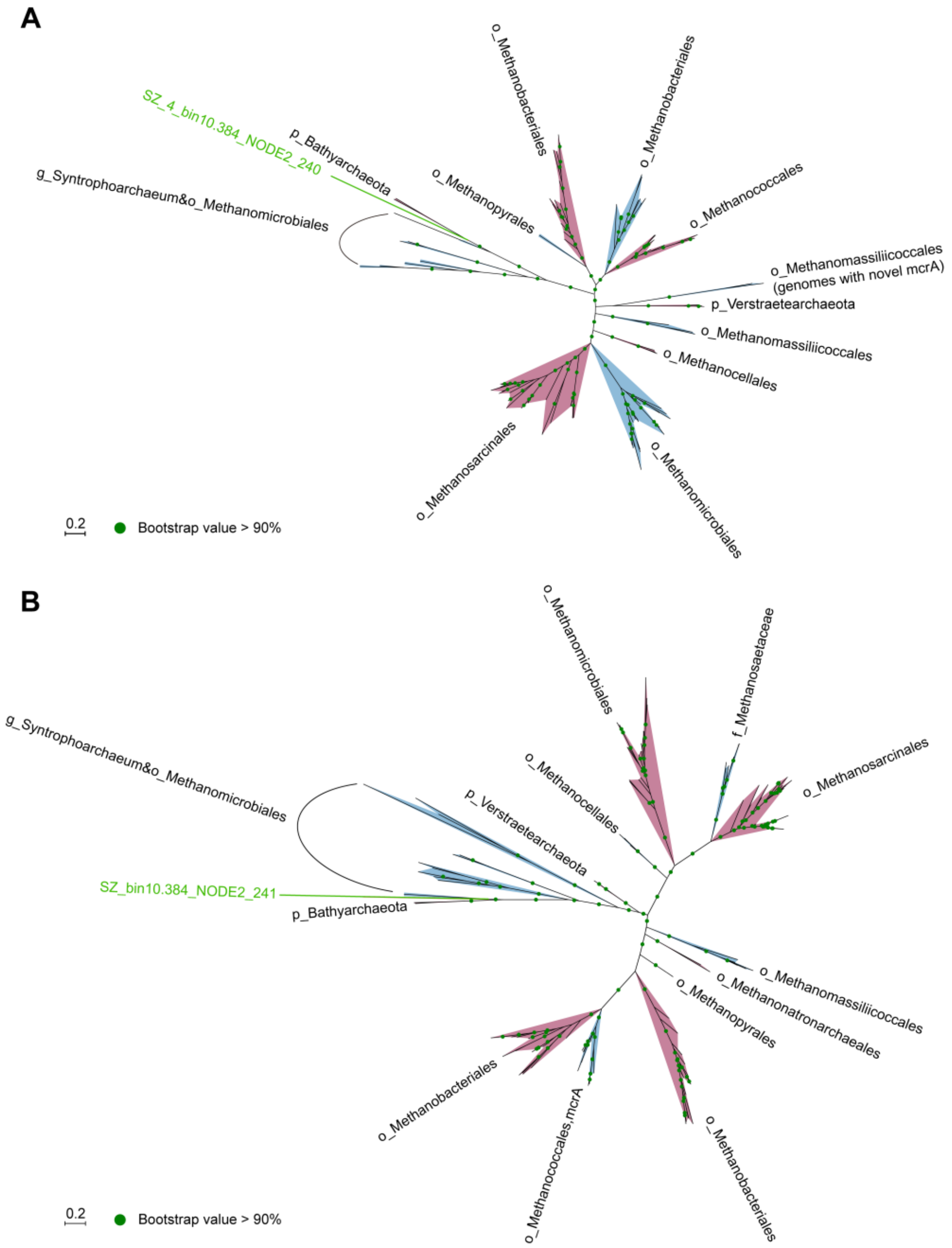


**Fig. S15. Protein trees of Asgard archaea (A) McrB and (B) McrG.** Asgard archaea McrA gene obtained in this study is marked with green line. The maximum-likelihood trees were inferred using IQ-TREE tree with model LG+F+I+G4 and parameter "-bb 1000". The background-colour (i.e., red and blue) is added to differentiate adjacent groups.

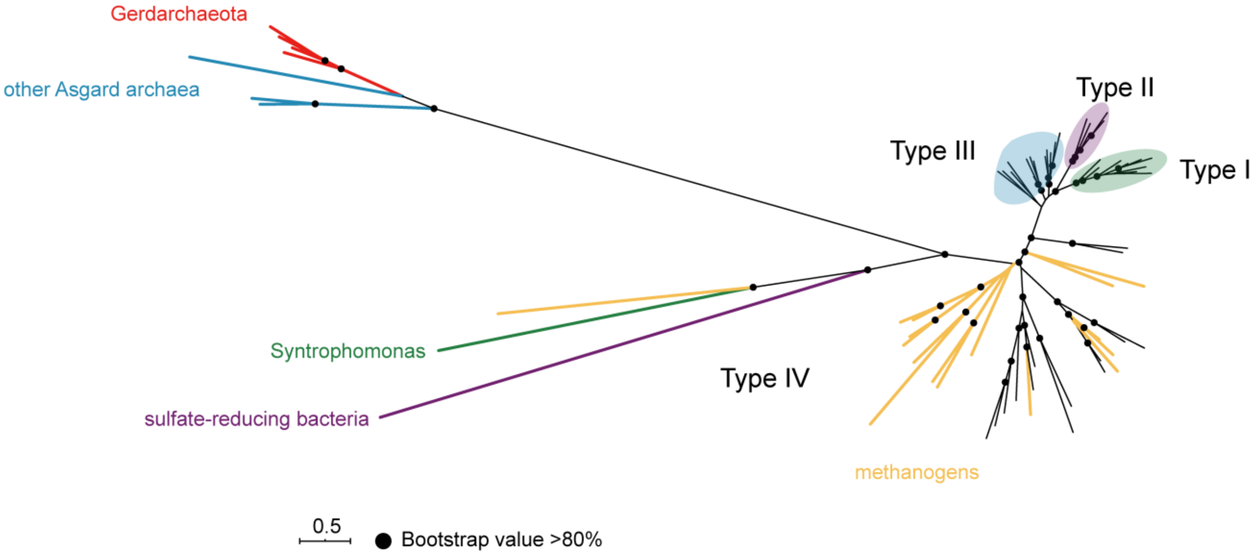


**Fig. S16. Phylogenetic position of the Gerdarchaeota *nifH*.** The unrooted maximum-likelihood tree was obtained using IQ-TREE software with the mixture mode 'LG+G4'. Asgard archaea *nifH* includes the novel ones obtained through BLASTN search against NCBI nr database, and those from NCBI database. Reference sequences were obtained from a previous study[95].

**References**


73. Quast, C. *et al.* The SILVA ribosomal RNA gene database project: improved data processing and web-based tools. *Nucleic Acids Res.* **41**, D590 (2012).
74. Kans, J. Entrez direct: E-utilities on the UNIX command line. (2017).
75. Edgar, R. C. Search and clustering orders of magnitude faster than BLAST. *Bioinformatics* **26**, 2460-2461 (2010).
76. Karst, S. M. *et al.* Retrieval of a million high-quality, full-length microbial 16S and 18S rRNA gene sequences without primer bias. *Nat. Biotechnol.* **36**, 190-195 (2018).
77. Ludwig, W. *et al.* ARB: a software environment for sequence data. *Nucleic Acids Res.* **32**, 1363-1371 (2004).
78. Konstantinidis, K. T., Rosselló-Móra, R. & Amann, R. Uncultivated microbes in need of their own taxonomy. *ISME J.* **11**, 2399 (2017).
79. Letunic, I. & Bork, P. Interactive Tree Of Life (iTOL): an online tool for phylogenetic tree display and annotation. *Bioinformatics* **23**, 127-128 (2006).
80. Berman, H. M. *et al.* The protein data bank. *Nucleic Acids Res.* **28**, 235-242 (2000).
81. Eswar, N. *et al.* Comparative protein structure modeling using Modeller. *Curr. Protoc. Bioinformatics* **15**, 5.6. 1-5.6. 30 (2006).
82. Morris, G. M. *et al.* AutoDock4 and AutoDockTools4: Automated docking with selective receptor flexibility. *J. Comput. Chem.* **30**, 2785-2791 (2009).
83. Morris, G. M. *et al.* Automated docking using a Lamarckian genetic algorithm and an empirical binding free energy function. *J. Comput. Chem.* **19**, 1639-1662 (1998).
84. Van Der Spoel, D. *et al.* GROMACS: fast, flexible, and free. *J. Comput. Chem.* **26**, 1701-1718 (2005).
85. Lindorff-Larsen, K. *et al.* Improved side-chain torsion potentials for the Amber ff99SB protein force field. *Proteins* **78**, 1950-1958 (2010).
86. Jorgensen, W. L. *et al.* Comparison of simple potential functions for simulating liquid water. *J. Chem. Phys.* **79**, 926-935 (1983).
87. Darden, T., York, D. & Pedersen, L. Particle mesh Ewald: An N⋅log (N) method for Ewald sums in large systems. *J. Chem. Phys.* **98**, 10089-10092 (1993).
88. Zhu, Q., Kosoy, M. & Dittmar, K. HGTector: an automated method facilitating genome-wide discovery of putative horizontal gene transfers. *BMC genomics* **15**, 717 (2014).
89. Steinegger, M. & Söding, J. MMseqs2 enables sensitive protein sequence searching for the analysis of massive data sets. *Nat. Biotechnol.* **35**, 1026-1028 (2017).
90. Yarza, P. *et al.* Uniting the classification of cultured and uncultured bacteria and archaea using 16S rRNA gene sequences. *Nat. Rev. Microbiol.* **12**, 635-645 (2014).
91. Eme, L. *et al.* Archaea and the origin of eukaryotes. *Nat. Rev. Microbiol.* **15**, 711-723 (2017).
92. Laso-Pérez, R. *et al.* Thermophilic archaea activate butane via alkyl-coenzyme M formation. *Nature* **539**, 396-401 (2016).
93. Chen, S.-C. *et al.* Anaerobic oxidation of ethane by archaea from a marine hydrocarbon seep. *Nature* **568**, 108-111 (2019).
94. Boyd, E. S., Hamilton, T. L. & Peters, J. W. An alternative path for the evolution of biological nitrogen fixation. *Front. Microbiol.* **2**, 205 (2011).
95. Gaby, J. C. & Buckley, D. H. A comprehensive aligned nifH gene database: a multipurpose tool for studies of nitrogen-fixing bacteria. *Database* **2014**, (2014).

96. Magnabosco, C. *et al.* A metagenomic window into carbon metabolism at 3 km depth in Precambrian continental crust. *ISME J.* **10**, 730-741 (2016).
97. D'hondt, S. *et al.* Distributions of microbial activities in deep subseafloor sediments. *Science* **306**, 2216-2221 (2004).
98. Søndergaard, D., Pedersen, C. N. S. & Greening, C. HydDB: A web tool for hydrogenase classification and analysis. *Sci. Rep.* **6**, 34212 (2016).
